# TDP-43 Sustains Satellite Cells to Maintain and Regenerate Skeletal Muscle

**DOI:** 10.64898/2026.05.18.725568

**Authors:** Theodore E. Ewachiw, Tenaya K. Vallery, Shloka Dhar, Hayden Clarkson, Tiffany Elston, Haileigh Gay, Bradley B. Olwin

## Abstract

Skeletal muscle satellite cells, residing between the myofiber plasma membrane and the surrounding basement membrane, maintain and repair skeletal muscle throughout life. Typically quiescent, satellite cells can transition into a reversible alert state (G_Alert_) that primes them for rapid activation to maintain or repair muscle. From G_Alert_, SCs can either re-enter quiescence or commit to the cell cycle, expand, and differentiate to fuse with existing regenerating myofibers. Exit from quiescence requires extensive post-transcriptional remodeling, including changes in RNA processing and RNA-binding protein activity. We show that TDP-43, an RNA binding protein, is essential for SC maintenance and muscle repair. Conditional deletion of TDP-43 in SCs caused a consistent and progressive loss of G_Alert_ SCs even in uninjured muscle, leading to depletion of the SC pool. TDP-43 haploinsufficiency was sufficient to impair SC maintenance, indicating that both alleles are required. Integrative analysis suggests that TDP-43 supports expression of stress response-associated transcripts during the quiescent-to-G_Alert_ transition, and that failure to mount this response contributes to SC apoptosis. Thus, we identified TDP-43 as a critical regulator of satellite cell survival as satellite cells activate and establish a TDP-43 requirement for maintaining and repairing skeletal muscle.

## Introduction

Skeletal muscle requires routine maintenance and repair by resident muscle stem cells or satellite cells (SCs).^1–4^ SCs occupy a specialized niche between the myofiber plasma membrane and the surrounding basement membrane and are typically quiescent. In response to physiological demand or injury, SCs transition into an alert state (G_Alert_) that primes them for rapid activation.^5,6^ From G_Alert_, SCs either re-acquire quiescence or enter G1, committing to DNA synthesis, terminally differentiate and fuse with existing myofibers, or form new myofibers.^5,6^ Although G_Alert_ accelerates cell cycle entry, sustained displacement from quiescence compromises long-term stem cell maintenance. SCs forced into G_Alert_ display impaired self-renewal and progressively fail to replenish the quiescent (G_0_) pool.^7^ Consistent with this, mice possessing constitutively G_Alert_ SCs exhibit early-life muscle hypertrophy but progressively deplete the quiescent SC pool during aging.^7,8^

Transitioning from quiescence to G_Alert_ and into the cell cycle requires extensive cell growth, organelle replication, and major transcriptomic and epigenetic remodeling.^5,9–14^ Among the most prominent events accompanying SC activation is altered expression and activity of RNA-binding proteins, underscoring the importance of post-transcriptional regulation during early cell fate decisions.^15,16^ For example, MyoD protein accumulates rapidly in activated SCs as MyoD1 mRNAs are regulated post-transcriptionally by the TTP family of RNA binding proteins,^17^ by Staufen1 mediated control of MyoD1 translation,^18^ and DEK dependent regulation of MyoD1 mRNA splicing.^19^ MicroRNAs^20^ and dynamic changes in translation^21,22^ regulate SC quiescence, highlighting the complexity of the post-transcriptional mechanisms involved preserving quiescence and enabling activation. RNA binding proteins with broad expression exert context-specific effects in SCs. For example AUF-1, an RNA decay enzyme, influences myogenic cell fate by promoting decay of MMP9 and altering the SC niche.^23^ In addition, HNRNPA2B1, a universally expressed RNA binding protein involved in splicing, uniquely disrupts cell fate transitions in myogenic cells when ablated in SCs,^24^ while CRISPR-Cas9-mediated knock out of TAR DNA-binding protein 43 (TDP-43; encoded by Tardbp) in myogenic C2C12 cells prevents cell growth and differentiation, promoting cell death.^25^

We conditionally depleted TDP-43 in SCs causing a consistent and progressive loss of SCs even in the absence of injury. Furthermore, TDP-43 ablated SCs failed to regenerate muscle upon injury. Using pseudotime analysis of single-cell RNA sequencing datasets, we identified stress response-associated gene programs that are induced as SCs transition from quiescence to cell cycle entry. Our data support a model in which TDP-43 is required for SC survival during the quiescence-to-G_Alert_ transition; in its absence, SCs undergo apoptosis and the stem cell pool collapses. Together, these findings establish TDP-43 as an essential regulator of early SC fate transitions required to maintain and repair skeletal muscle.

## Results

### SCs require TDP-43 to regenerate skeletal muscle

CRISPR-mediated knock out of TDP-43 in C2C12 myoblasts induces cell death and prevents terminal differentiation, suggesting that TDP-43 is required to transit cell fates and for myogenic cell survival.^25^ We assessed whether SCs require TDP-43 *in vivo* by crossing *Pax7^CreERT2^* mice with *Tardbp^loxp/loxp^*mice to obtain *Pax7^CreERT2^*;*Tardbp^loxp/loxp^*inducible SC-specific knockout mice (KO mice). Since *Tardbp* and *Pax7* are located in close proximity to each other on chromosome 4, recombination between the two nearby loci was verified by genotyping (Fig. S1A-C). Recombination efficiency was assessed by quantifying TDP-43 immunoreactivity in SC nuclei from serial tibialis anterior (TA) muscle sections and by qPCR of isolated SCs (Fig. S1D-G). TDP-43 expression was significantly reduced in SCs isolated from KO mice and from *Pax7^CreERT2^*;*Tardbp^+/loxp^*heterozygous mice (HET), compared to *Pax7^CreERT2^*;*Tardbp^+/+^* controls (WT) (Fig. S1G). Accompanying the decrease in TDP-43 expression, we measured a 2-fold decline of TDP-43 transcript levels in HET SCs and 5.5-fold decline in KO SCs (Fig. S1H), was a significant loss of SC numbers in KO muscle. Of the remaining SCs in KO muscle, 25% were immunoreactive for TDP-43 compared to ≥ 90% TDP-43+ SCs in WT mice (Fig. S1I).

We injured TA muscles with BaCl_2_ and analyzed tissue at 10 days post-injury (10 dpi) to test if TDP-43 is required for SCs to regenerate muscle. Muscles in KO mice contained few, if any, regenerating myofibers, whereas muscles in HET mice regenerated comparably to WT controls (Fig. 1A). Few SCs are detected in KO mouse muscle at 10dpi, while HET and WT mice possess similar SC numbers (Fig. 1B, C). Consistent with the loss of SCs, only 10% of myofibers in KO muscle at 10 dpi possess centrally located nuclei (Fig. 1D) and only 10% of the TA cross-sectional area regenerated at 10dpi (Fig. 1E). Fusion of SCs lacking TDP-43 in regenerating muscle was assessed by crossing KO mice with *LSL:tdTomato* reporter mice, generating *Pax7^CreERT2^*;*Tardbp^loxp/loxp^;LSL:tdTomato* mice (KO^tdT^), *Pax7^CreERT2^*;*Tardbp^+/loxp^;LSL:tdTomato* mice (HET^tdT^) and *Pax7^CreERT2^*;*Tardbp^+/+^;LSL:tdTomato* mice (WT^tdT^; Fig. S1D). Following tamoxifen administration, SCs will be lineage marked with tdTomato and fusion of tdTomato lineage-marked SCs will result in tdTomato+ myofibers. At 28 dpi, nearly all myofibers in WT^tdT^ muscle were tdTomato+, whereas KO^tdT^ muscles contained very few tdTomato+ myofibers (Fig. 1F). HET^tdT^ muscle displayed an intermediate phenotype (Fig. 1F). Because WT^tdT^ and HET^tdT^ mice contained comparable numbers of SCs at 28 dpi (Fig. 1G), the reduction in tdTomato+ myofibers in HET^tdT^ muscle reflects impaired fusion rather than differences in SC abundance. In KO^tdT^ muscle, the near absence of tdTomato+ myofibers mirrored the profound (95%) loss of SCs (Fig. 1F, G). The small fraction of SCs remaining in KO^tdT^ muscle at 28 dpi were uniformly TDP-43+ (Fig. 1H) and likely escaped recombination.

**Fig. 1.**
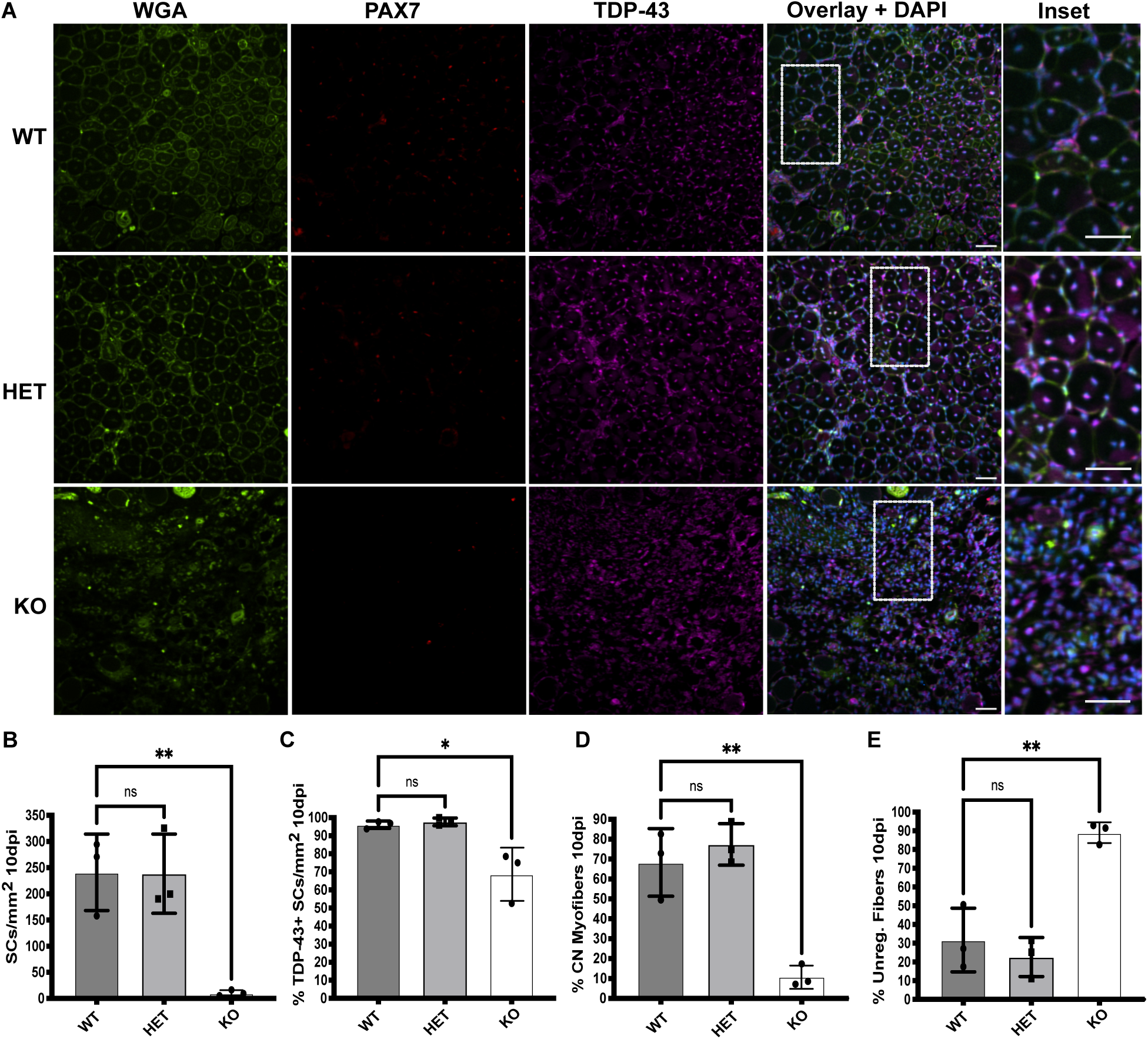
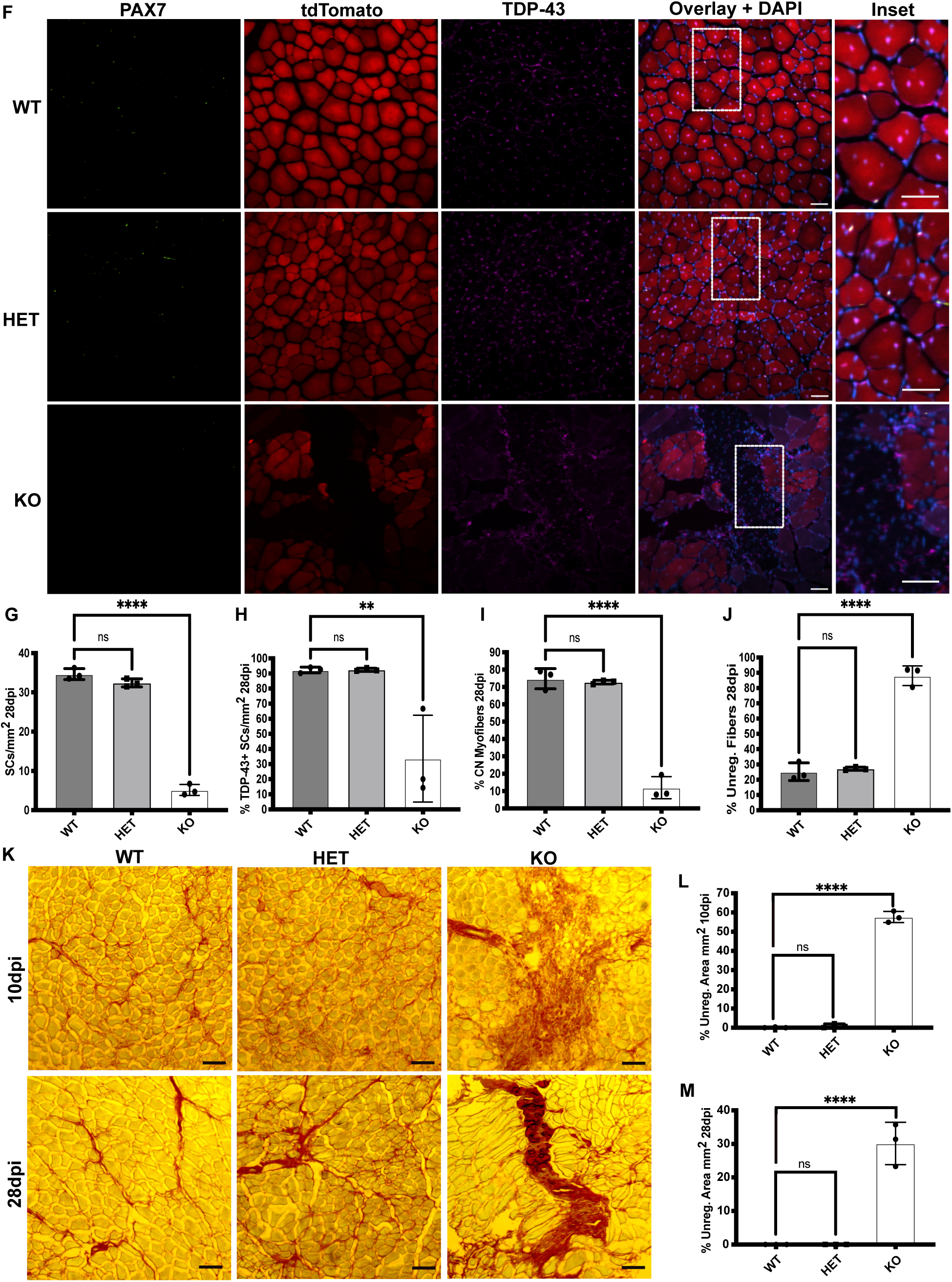
SCs require TDP-43 to Regenerate Muscle. **(A)** Representative images of regenerating TA muscle from Pax7^CreERT2^;TDP-43^flox/wt^ and Pax7^CreERT2^;TDP-43^flox/flox^ mice at 10dpi with myofiber boundaries identified by fluorescent WGA, SCs identified by Pax7 immunoreactivity and TDP-43 protein assessed by immunoreactivty. Scale bar: 50µm. The numbers of **(B)** SCs, **(C)** TDP-43 immunoreactive SCs,(**D**) **m**yofibers with centrally located nuclei, and **(E)** Percent unregenerated fibers were quantified at 10dpi. **(F)** Representative images regenerating TA muscle at 28dpi from Pax7^CreERT2^;TDP-43^wt/wt^;LSL:tdTomato (WT^tdT^), Pax7^CreERT2^;TDP-43^flox/wt^;LSL:tdTomato (HET^tdT^) and Pax7^CreERT2^;TDP-43^flox/flox^;LSL:tdTomato (KO^tdT^) mice with SCs identified by Pax7 immunoreactivity, recombined SCs and myofibers possessing recombined SCs identified by tdTomato flouresecence and nuclei identified by DAPI fluorescence. Scale bar: 50µm. The numbers of **(G)** SCs per mm^2^, **(H)** TDP-43+ SCs, **(I)** myofibers with centrally located nuclei and, **(J)** unregenerated myofibers were quantified at 28dpi. **(K)** Representative Picrosirius Red images of regenerating TA muscle at 10dpi and 28dpi from Pax7^CreERT2^;TDP-43^flox/wt^ and Pax7^CreERT2^;TDP-43^flox/flox^ mice. Scale bar: 100µm. The **(L)** percent unregenerated TA muscle area at 10dpi and **(M)** percent unregenerated TA muscle area at 28dpi was quantified. n = 3 mice per genotype. For all statistical analysis, Ordinary one-way ANOVA was used to compare groups. * P = < 0.05, ** P = < 0.003, *** P = 0.0003, **** P = < 0.0001. See also Figure S1.

Regenerative outcomes were similarly impaired in KO^tdT^ muscle at 28 dpi, with few centrally nucleated myofibers (Fig. 1I) and persistent failure of tissue regeneration (Fig. 1J). Fibrosis assayed by Picrosirius Red staining was minimal in WT and HET muscle at 10dpi and 28dpi (Fig. 1K-M). In stark contrast, KO muscles exhibited substantial fibrosis, comprising 30% of the TA at 10dpi and 50% by 28 dpi (Fig. 1K-M). TDP-43 is thus essential for SC survival and regenerative myogenesis, and TDP-43-null SCs are lost prior to contributing to myofiber repair.

### SCs fail to maintain skeletal muscle and die following TDP-43 deletion *in vivo* and *in vitro*

TDP-43 may be necessary for muscle maintenance since muscles in KO mice fail to regenerate following injury and SC are lost prior to fusion during muscle repair. We quantified tdTomato+ SCs and tdTomato+ myofibers in uninjured WT^tdT^, HET^tdT^, and KO^tdT^ mice 28 days post tamoxifen injection (Fig. 2A) where fewer tdTomato+ myofibers were evident in HET^tdT^ and KO^tdT^ muscle compared to WT^tdT^ muscle (Fig. 2B). Approximately 35% of myofibers in WT^tdT^ mice were tdTomato positive at 28 days post-recombination (Fig. 2A)^1,27^ with few tdTomato+ myofibers visible in KO^tdT^ mice (Fig. 2B). SC abundance was reduced 1.7-fold (Fig. 2C), with a near complete loss of TDP-43+ SCs (Fig. S2A, B).

**Fig. 2.**
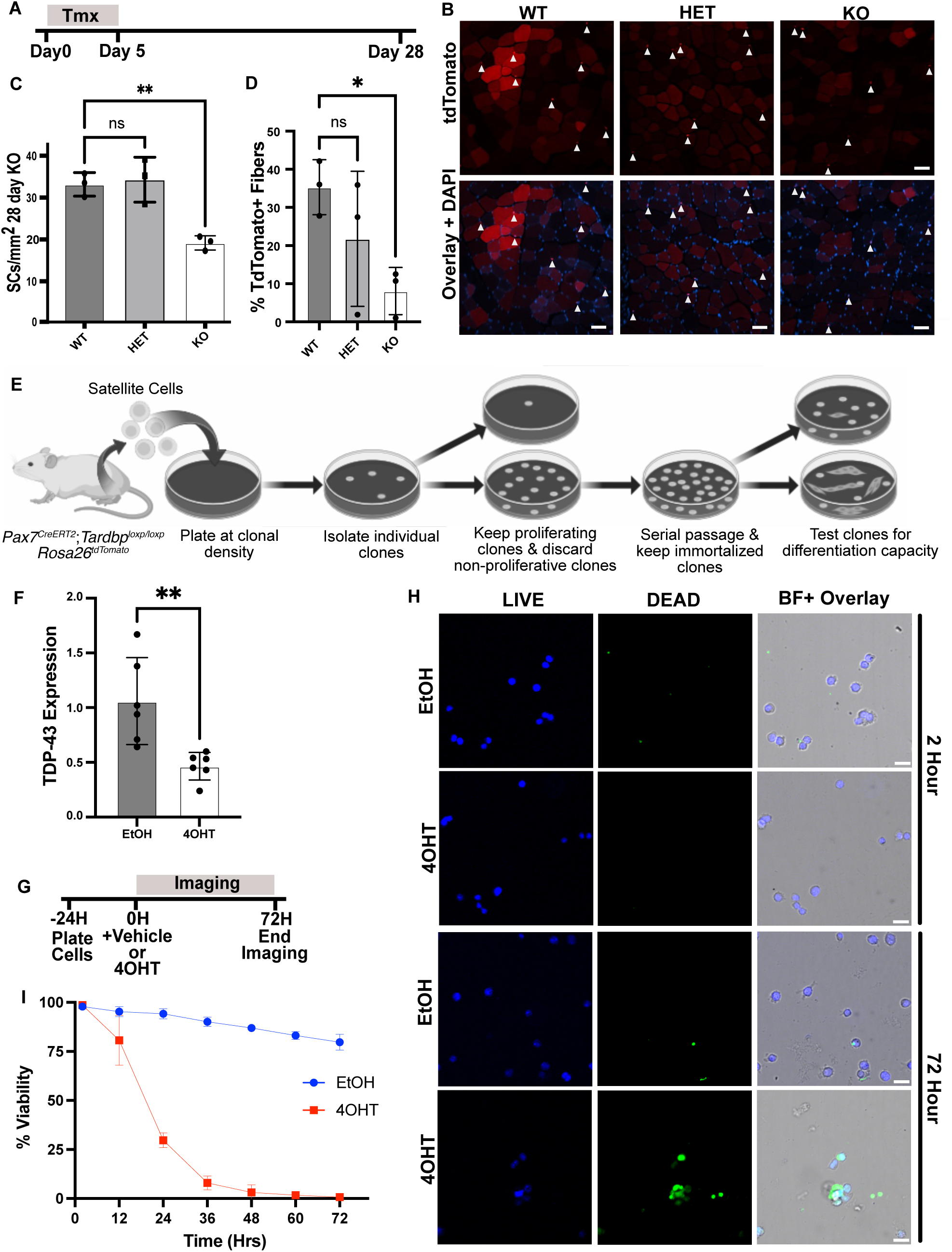
TDP-43 null SCs are unable to maintain muscle. **(A)** Timeline for TDP-43 ablation. **(B)** Representative images of TA muscles from WT^tdT^, HET^tdT^ and KO^tdT^ mice 28-day post recombination (solid arrowheads identify SCs). Scale bar: 50µm. **(C)** SC number and percent tdTomato+ myofibers in mouse TA muscle sections quantified at 28d post recombination for WT^tdT^, HET^tdT^ and KO^tdT^ mice. n = 3 mice per genotype. One-way ANOVA ** P = < 0.004. **(D)** TtdTomato fluorescent intensity distribution reflecting fusion of recombined SCs in WT^tdT^, HET^tdT^ and KO^tdT^ mice 28-day post recombination (> 480 myofibers quantified per condition from n = 3 mice). One-way ANOVA * P = < 0.05. **(E)** Schematic for producing an immortalized cell line from unrecombined KO mice. **(F)** TDP-43 quantified by qRT-PCR in recombined and unrecombined KO cells 72 following vehicle or 4OHT treatment. n = 6 independent experiments. Unpaired, two-tailed Students t-test ** P = < 0.006. Data are mean ± s.d. **(G)** Schematic for experimental time line of live cell imaging in KO cells. **(H)** Representative images from live cell microscopy of unrecombined and recombined KO cells assayed for cell integrity with ReadyProbes Cell Viability kit. Scale bar = 25µm. **(I)** Percent viability of unrecombined or recombined KO cells for 72-hour following TDP-43 ablation. n = 3 biological independent experiments. Two-way ANOVA. 12 h P = 0.0017, h 24 through 72 P = < 0.0001. See also Figure S2.

TdTomato+ myofibers were reduced 4-fold relative to WT^tdT^ (Fig. 2D) where the magnitude of tdTomato+ myofiber loss exceeded the reduction in SC number, consistent with the interpretation that TDP-43 null SCs die prior to fusion into myofibers.

In HET^tdT^ mice SC numbers were comparable to WT^tdT^ (Fig. 2C) with similar numbers of TDP-43+ SCs (Fig. S2B). However, tdTomato+ myofibers were reduced by 2-fold in HET^tdT^ mice (Fig. 2D), demonstrating TDP-43 haploinsufficiency for SC contribution to myofiber maintenance. The decrease in fusion was consistent with reduced centrally located myonuclei in HET^tdT^ and KO^tdT^ muscle at 28 days post-recombination (Fig. S2D, E). Despite reduced fusion, myofiber morphometry was not significantly different between mouse cohorts (Figure S2C) as TDP-43 is present in the majority of myonuclei and only fusing SCs lack TDP-43.

To determine the timing of SC sensitivity to TDP-43 loss, we generated an immortalized SC line from KO^tdT^ mice (Fig. 2E). Treatment with 4-hydroxytamoxifen (4OHT) reduced TDP-43 expression by >2-fold within 72 hours compared to vehicle-treated controls (Fig. 2F). Live-cell imaging enabled us to quantify SC number and viability in real time using a cell-impermeant dye that labels cells with compromised membrane integrity (Fig. 2G, H). As early as 12 hours after 4OHT treatment, ∼25% of KO^tdT^ cells were dye-positive, increasing to ∼75% by 24 hours (Fig. 2H, I). By 72 hours, nearly all 4OHT-treated KO^tdT^ cells are dye-positive, compared to ∼10% of vehicle treated controls (Fig. 2H, I). Consistent with this rapid loss of viability, growth curves revealed no proliferative expansion following recombination (Fig. S2F, G). Therefore, SCs die rapidly following TDP-43 deletion, prior to completing the first cell cycle.

### TDP-43 loss induces SC apoptosis

Given the rapid SC loss following TDP-43 deletion, we tested whether SCs die via apoptosis, necrosis, or anoikis. TA muscles were harvested 10 days after recombination and analyzed for markers of cell death (Fig. 3A). Cytochrome-c and cleaved-PARP1 immunoreactivity revealed that >50% of SCs in KO muscle were apoptotic (Fig. 3B, C), increasing 6.7-fold and 5.1-fold increases, respectively, compared to WT controls (Fig. 3D, E). HET SCs exhibited a ∼2-fold increase in cleaved-PARP1 immunoreactivity relative to WT SCs, although cytochrome-c immunodetection was not significantly altered (Fig. 3B, C).

**Fig. 3.**
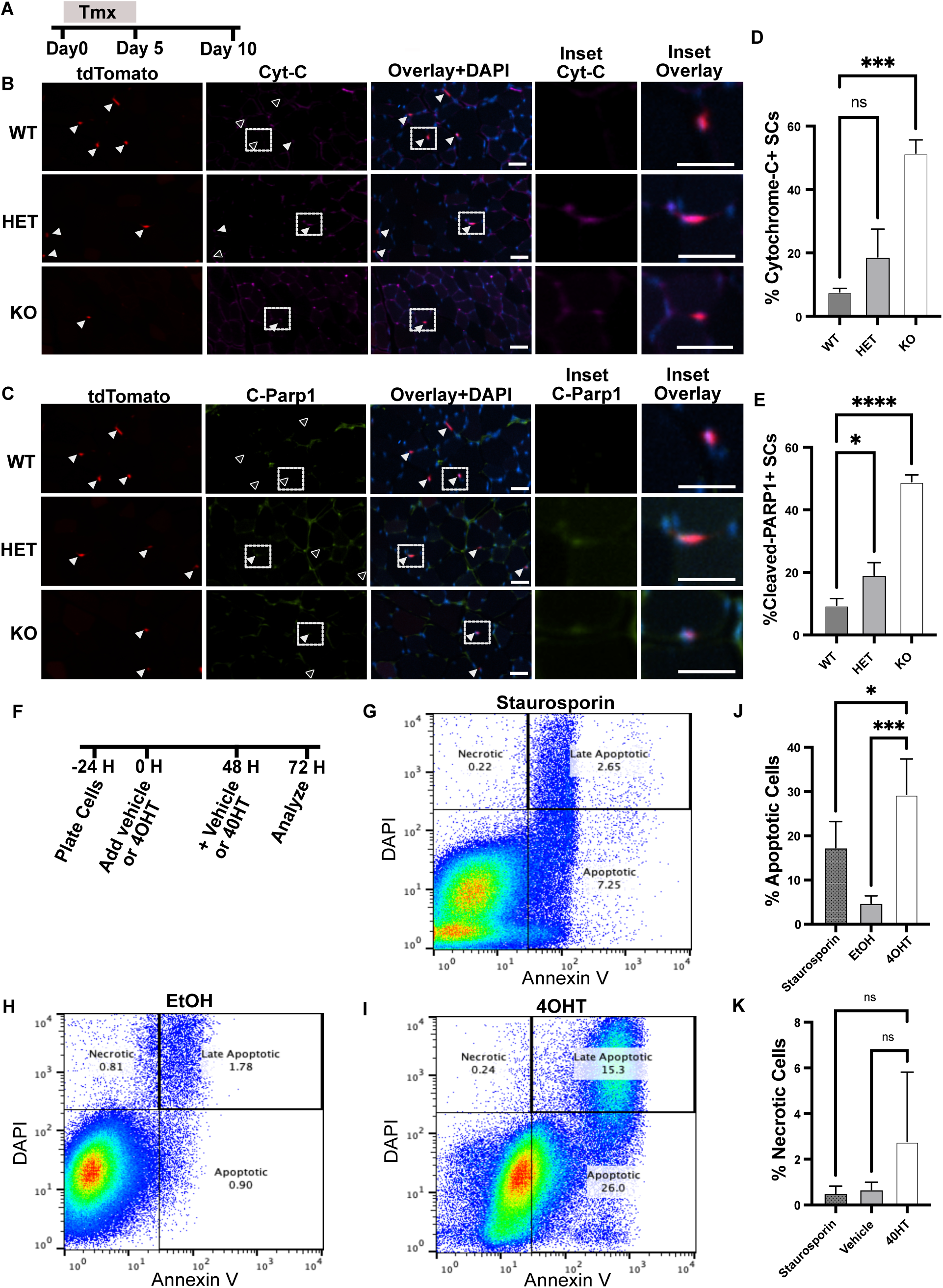
SCs apoptose following TDP-43 deletion. **(A)** Timeline for tamoxifen injections and tissue harvest. **(B, C)** Representative images of TA muscles from WT^tdT^, HET^tdT^ and KO^tdT^ mice 10 days post recombination assayed for cytochrome C immunoreactivity. Solid arrowheads identify SCs positive for tdTomato, cytochrome-C and both (**B**), or positive for tdTomato, cleaved-PARP1 or both (**C**). Clear arrowheads identify SCs that are not immunoreactive for either cytochrome-C or for cleaved-PARP1. Scale bar: 50µm. **(D)** SCs quantified for cytochrome-c immunoreactivity and, **(E)** cleaved-PARP1 immunoreactivity from WT^tdT^, HET^tdT^ and KO^tdT^ mice 10 days post recombination. n = 3 mice of each genotype. **(F)** Schematic of cell culture and treatment preceding flow cytometry for WT cells. **(G-I)** Flow cytometry gating for WT SCs analyzed at 72-hours post treatment. KO cells were treated with 1µM of staurosporin for 6 hours prior to analysis as a positive control. A minimum of 6 x 10^5^ cells were analyzed in each condition. KO cells from three independent flow cytometry analyses were quantified for **(J)** apoptosis and **(K)** necrosis. n = 3 biological independent experiments. For all statistical analyses (D, E, K, J), Ordinary one-way ANOVA was used. * P = < 0.05, *** P = < 0.0004, **** P = < 0.0001.

To validate apoptosis *in vitro,* we assessed recombined KO^tdT^ SCs by flow cytometry using Annexin V and DAPI staining, with staurosporine-treated cells used as a positive control for apoptosis (Fig. 3F). At 72 hours following 4OHT treatment, Annexin V+/DAPI+ staining increased substantially, indicating robust apoptosis upon TDP-43 loss (Fig. 3G-I). A ∼6-fold increase in apoptosis occurred in recombined KO SCs relative to vehicle-treated controls (Fig. 3J), exceeding the ∼2-fold increase in staurosporine-treated unrecombined controls (Fig. 3J). In contrast, necrosis was not significantly increased in any condition (Fig. 3K), consistent with the conclusion that loss of TDP-43 triggers rapid apoptotic death of SCs.

### Identifying SC cell fate transitions

To gain mechanistic insight into why SCs undergo apoptosis following TDP-43 loss, we examined TDP-43 RNA-binding targets using enhanced ultraviolet crosslinking and immunoprecipitation (eCLIP) performed in myoblasts and compared these targets to those previously identified in myotubes.^25^ We identified 106 transcripts that were significantly enriched in myoblasts relative to myotubes (Fig. 4A). In myoblasts, TDP-43 binding occurred predominantly within exons of protein-coding transcripts (Fig. 4B), consistent with a role in regulating mRNA processing and stability. Gene ontology (GO) analysis of myoblasts-enriched targets revealed strong enrichment for biological processes associated with nuclear transport and cellular stress responses (Fig. 4C), and the top terms included pathways related to nuclear export, plasma membrane repair, stem cell maintenance, and regulation of cell fate decisions (Fig. 4D). Reactome pathway analysis further highlighted stress response networks, including protein folding and regulation of apoptosis, as the most significantly enriched pathways among myoblast targets (Fig. 4E). These functional categories are consistent with the physiological demands placed on SCs as they exit quiescence and progress toward cell cycle entry.^5,9–14^

**Fig. 4.**
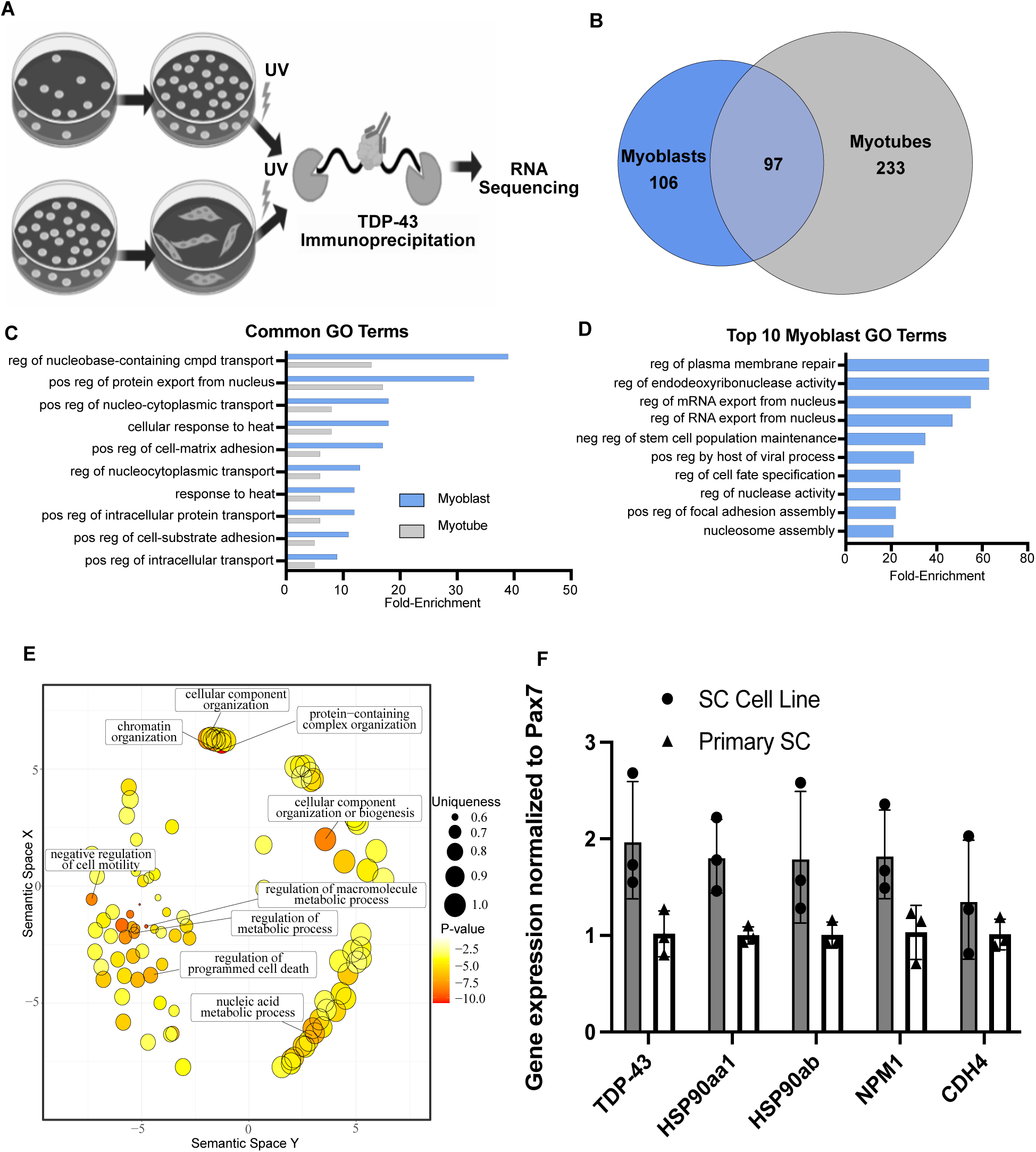
TDP-43 induces SC stress response transcripts. **(A)** Schematic of TDP-43 eCLIP subsequent RNA sequencing in myoblasts and myotubes. **(B)** Venn Diagram depicting enriched TDP-43 targets in myoblasts and myotubes. **(C)** Top 10 common biological processes by fold enrichment of TDP-43 eCLIP targets by Panther GO Slim **(D)** Top 10 TDP-43 myoblast reactome targets by eCLIP. **(E)** Heat map of TDP-43 eCLIP myoblast cellular processes. (**F**) Expression of stress response TDP-43 eCLIP targets by qRT-PCR of RNA in isolated SCs from unrecombined KO mice and in an unrecombined KO SC cell line.

Extensive cell and organelle growth occurs prior to the initial cell division at 24 to 36 hours after SCs activate.^5,9,11,12,15^ Transitioning from a quiescent state to DNA synthesis is intrinsically stressful, and SCs from aged mice transition from G_0_ to the cell cycle slower than SCs from young mice, often apoptosing.^30^ Based on the enrichment of stress response and apoptosis-associated pathways among myoblast TDP-43 targets, TDP-43 may support SC survival by enabling stress response gene expression as SCs transition into the cell cycle. We selected four putative TDP-43 targets identified in the eCLIP dataset-HSP90aa1, HSP90ab, NPM1 and CDH4-and assessed their expression by qPCR in freshly isolated SCs and in the unrecombined KO SC cell line (Fig. 4F) where these transcripts were readily detectable, supporting their relevance in SC biology.

We integrated single cell RNA-Seq, SC-enriched scRNA-Seq, and single nuclear RNA-Seq data from uninjured TA muscles and regenerating muscles at 4dpi and 7dpi^31^ to define when these stress-associated transcripts are induced during regeneration. After integrating and clustering using Seurat, we identified myogenic populations corresponding to Pax7+ SCs (Fig. S5A), myogenin-expressing differentiating myoblasts (Fig. S5B) and Tmem38a expressing myonuclei (Fig. S5C; Fig. 5A). We then mapped expression of canonical stress-response genes onto UMAP projections, using Calcitonin receptor (Calcr) to identify quiescent SCs (Fig. 5C), Id3 to identify activated and proliferating SCs (Fig. 5D), and the putative TDP-43 targets Hsp90aa1, Hsp90ab1, Npm1, and Cdh4 (Fig. 5E-H). Across myogenic populations, expression of Hsp90aa1, Hsp90ab1, and Npm1 was low in quiescent SCs but increased following injury, consistent with an increase as SCs transition to proliferating myoblasts (Fig. 5I-N, S5). In contrast, Cdh4 decreased after injury, mirroring the decline in the calcitonin receptor (Calcr) e xpressed in quiescent SCs (Fig. 5I,N). At 4 dpi Id3, Hsp90aa1, Hsp90ab1, and Npm1 expression increased significantly (Fig. 5J-M), whereas by 7dpi only Npm1 remained significantly elevated (Fig. 5M). While Calcr declined and Id3 increased as expected, these changes did not strongly shift UMAP localization (Fig. 5I,J; Fig. S5D,E). In contrast, Hsp90aa1, Hsp90ab1, Npm1, and CDH4 showed broader distribution across clusters, consistent with dynamic expression across multiple myogenic states (Fig 5K-N, Fig. S5F-I)

**Fig. 5.**
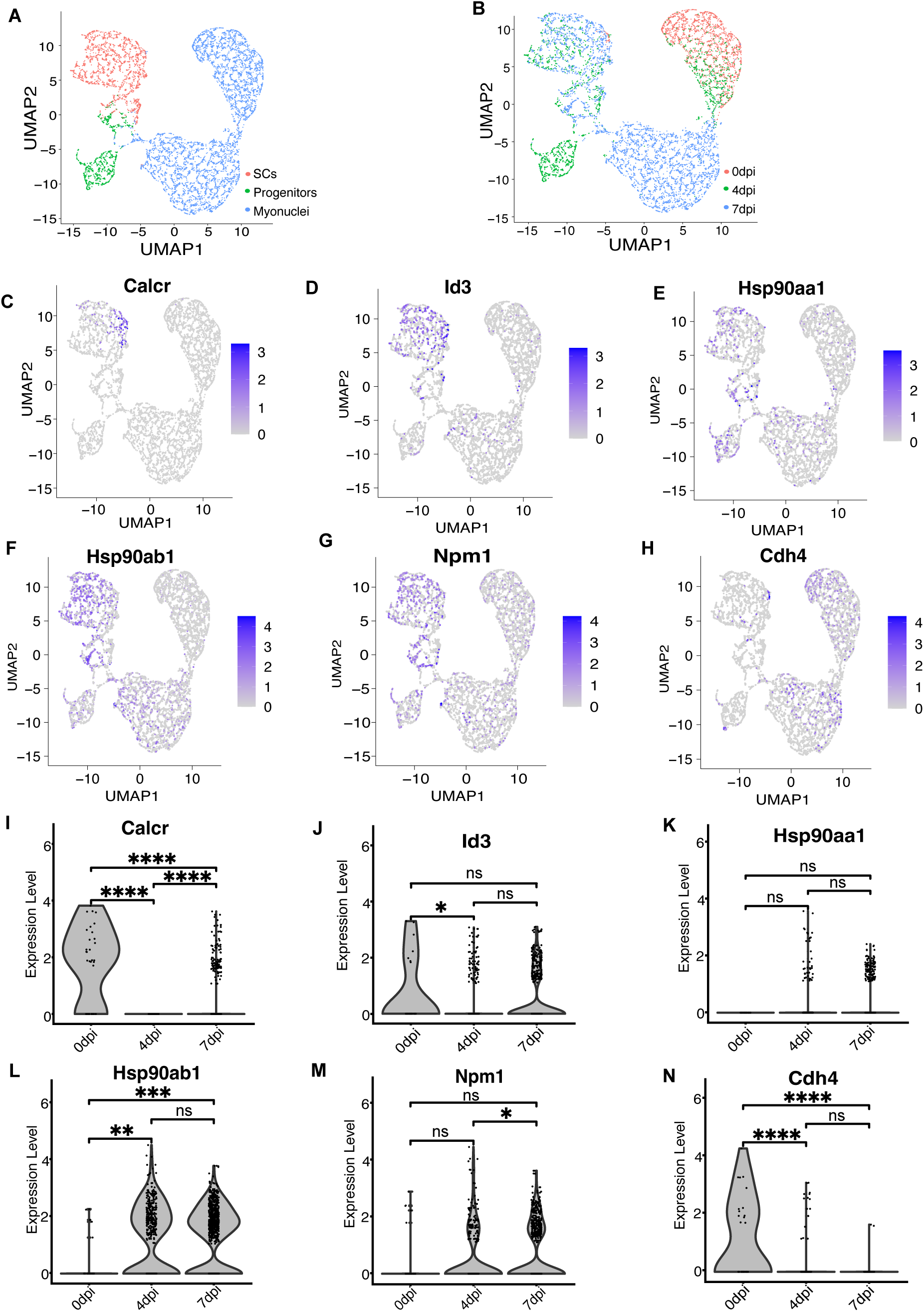
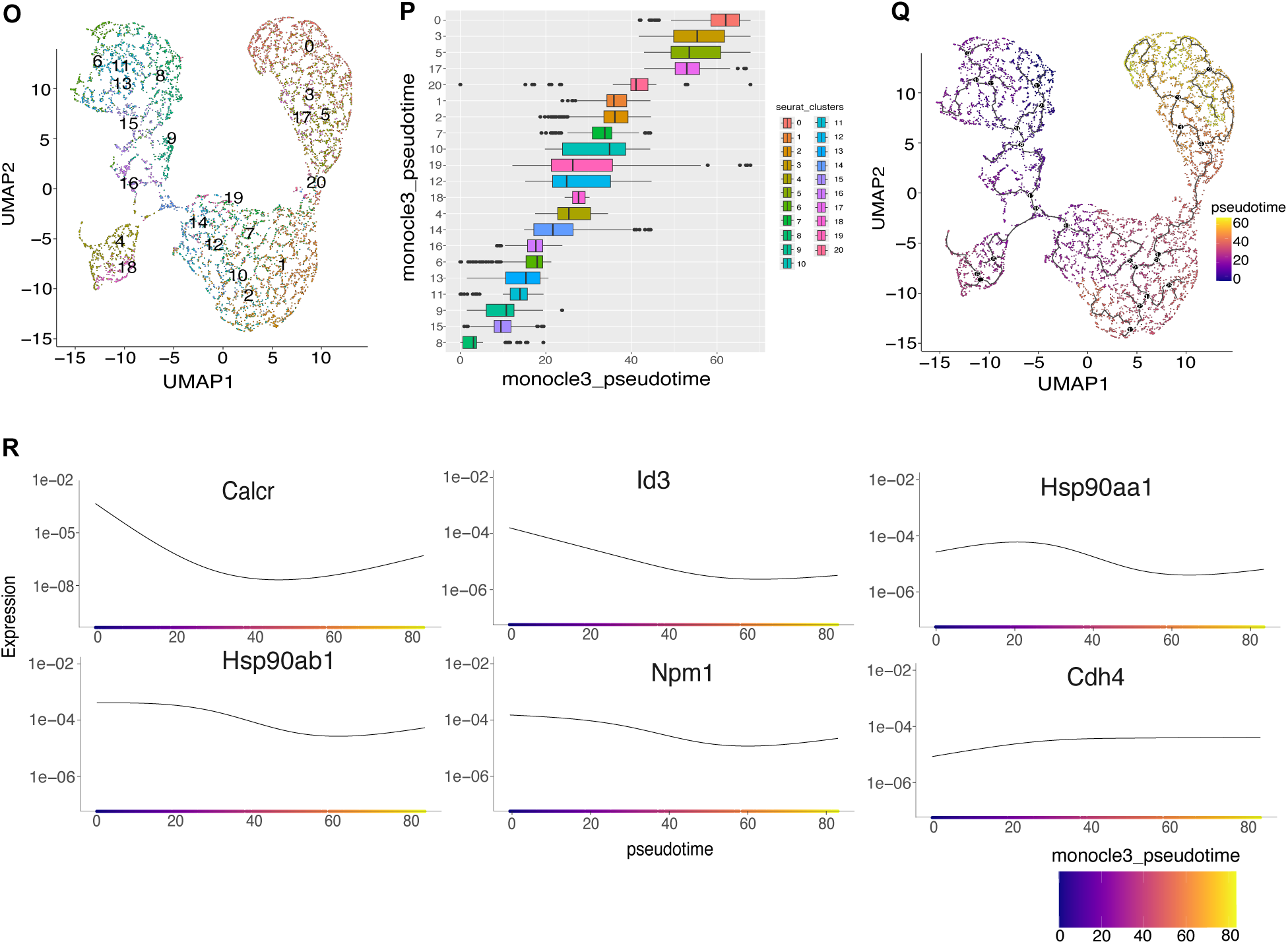
TDP-43 induced stress response transcripts are regulated as SCs change fate following an unjury. **(A)** UMAP clustering of nuclei from sequencing nuclei and single cells from skeletal muscle at 0, 4, and 7dpi. (n = 2 biological independent experiments).**(B)** UMAP clustering of nuclei from skeletal muscle colored to show predominant myogenic cell populations at 0, 4, and 7dpi. **(C, D** SC clusters identified quiescent SCs by Calcr expression (**C**), and by Id3 expression (**D**). **(E-H)** Subsets of myogenic nuclei that express cell-stress pathway genes identified as TDP-43 binding partners by eCLIP. **(I, J)** Violin plots of genes used to identify cell types from snRNA-seq at 0, 4, and 7dpi. * P = < 0.05, **** P = < 0.000006. **(K-N)** Violin plots for verified of verified TDP-43 target stress genes at 0, 4, and 7dpi. * P = < 0.05, ** P = < 0.006, *** P = < 0.0006, **** P = < 0.000006. **(O)** UMAP of clusters identified by PCA analysis revealing unique cell populations. **(P)** Monocle-inferred pseudotime clustering of myogenic cells in Q. **(Q)** Monocle-inferred pseudotime trajectory of myogenic nuclei colored by assigned pseuodotime value. **(R)** Pseudotime of myogenic cell population identifiers (ID3, Castor2, Calcr) and of TDP-43 targets identified by TDP-43 eCLIP (Hsp90aa1, Hsp90ab1, Cdh4, and Npm1). See also Figure S3.

We performed pseudotime trajectory analysis, anchoring the trajectory at cluster 8 and progressing toward cluster 0 (Fig. 5O-R) to better map gene expression with SC cell fate transitions. As expected, Calcr expression was highest at pseudotime 0, declined sharply through intermediate pseudotime, and modestly increased again later in the trajectory (Fig. 5R). In contrast, Hsp90aa1, Hsp90ab1and Npm1 were elevated early-to-mid trajectory (pseudotime 0 to 50/60), corresponding to the transition from quiescence into the cell cycle (Fig. 5R). Cdh4 showed a modest increase as SCs progressed from quiescence to proliferating myoblasts and differentiated myonuclei (Fig. 5R). Putative TDP-43-dependent stress response transcripts are induced at their highest levels as SCs exit quiescence and transit into the cell cycle and then terminally differentiate. The timing aligns with the rapid SC apoptosis observed upon TDP-43 deletion, supporting the idea that TDP-43 is required to sustain stress-response genes for SC survival as SCs transit cell fate changes. Because TDP-43-null SCs die rapidly after recombination (Fig. 2.H,I; Fig. S2D,E,H,I), we were unable to directly quantify these stress-response transcripts in KO SCs either immediately after recombination *in vivo* or following 4OHT-induced recombination *in vitro*.

### SCs are unable to transition from G_0_ to G_Alert_ following TDP-43 deletion

Ablating TDP-43 in cultured SCs or from SCs *in vivo* following a muscle injury induces rapid cell loss (Figs. 1-3) that appears to be caused by cells stress occurring upon transitioning cell fates. To determine which transition is most sensitive to TDP-43 loss, we focused on the reversible G_Alert_ state, which occurs prior to G1 entry and primes SCs for rapid cell cycle progression.^5^ SCs enter G_Alert_ not only in response to local injury, but also systemically; a unilateral muscle injury induces G_Alert_ in a subset of SCs in distant, uninjured muscles that returns to baseline levels by 28dpi.^5^ G_Alert_ SCs rapidly phosphorylate ribosomal protein S6 (pS6), providing a marker for quantifying G_alert_ SCs *in vivo*.^5^

To establish baseline levels of G_Alert_ SCs (QSCs) in the absence of injury, we quantified Pax7+ SCs and pS6 immunoreactivity in uninjured TA muscles from WT^tdT^, HET^tdT^ and KO^tdT^ mice at 10 and 28 days following recombination (Fig. 6A-E). At 10 days post-recombination, 1.6-fold fewer QSCs are present in KO^tdT^ muscle relative to WT^tdT^ (Fig 6B). However, pS6+ QSCs loss was substantially greater than SC loss as 5-fold fewer pS6+ QSCs are detected in KO^tdT^ mice compared with WT^tdT^ controls (Fig. 6B-D). By 28 days post-recombination, QSC numbers in KO^tdT^ muscle declined further (∼1.75-fold relative to WT^tdT^), with pS6+ QSCs reduced by ∼4-fold, similar to that at 10 dpi (Fig. 6C, E). Representative images from each genotype show Pax7+ QSCs and pS6 immunoreactivity at 10 and 28 days post-recombination, with filled arrowheads indicating pS6+ QSCs and open arrowheads indicating pS6-QSCs (Fig. 6D, E).

**Fig. 6.**
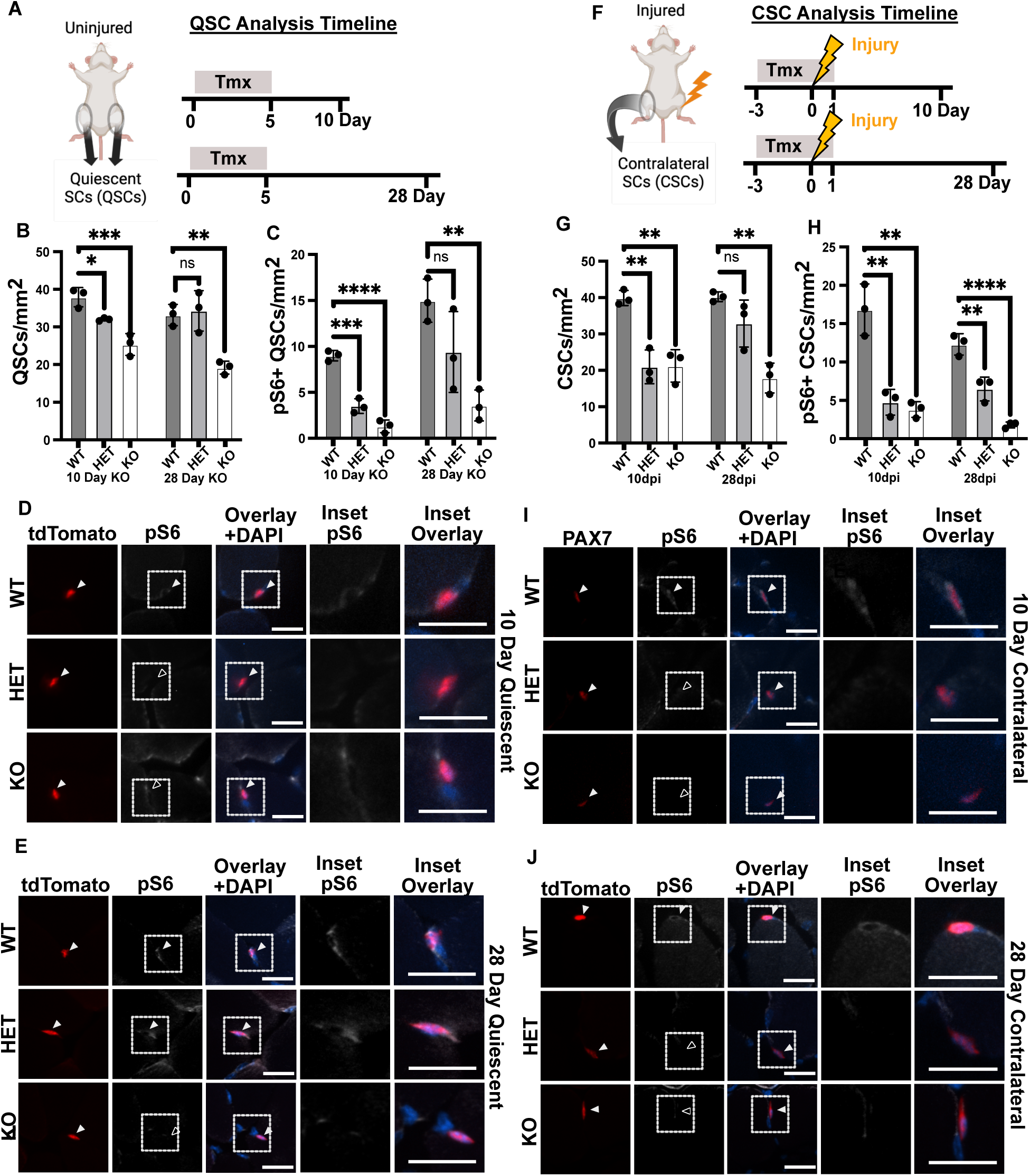
TDP-43 null SCs fail to enter G-Alert or maintain G_alert_. **(A)** Schematic for experimental timeline and QSC isolation from mice. **(B)** QSCs/mm^2^ quantified at 10d and 28d post tamoxifen injection in WT^tdT^, HET^tdT^ and KO^tdT^ mice. **(C)** CSCs/mm^2^ immunoreactive for pS6+ quantified as in (**B**). CSC timeline schematic for 10 and 28dpi TDP-43 KO. **(D)** Representative images from TA muscle 10d post tamoxifen injection identifying tdT+ QSCs from WT^tdT^ HET^tdT^ and KO^tdT^ mice where solid arrowheads indicate cells positive for tdTomato fluorescence or phospho-S6 immunoreactivity and open arrowheads indicating SCs negative for phospho-S6 immunoreactivity. Scale bar: 20µm. **(E)** Representative images from TA muscle 28d post tamoxifen injection identifying tdT+ QSCs immunoreactive for Pax7 and immunoreactive for pS6 in WT^tdT^ but negative for PS6 immunoreactivity in HET^tdT^ and KO^tdT^ mice with arrowheads as in (**D**). Scale bar: 20µm. **(F)** Schematic for injury, timeline and CSC isolation from contralateral TA muscle in mice. CSCs/mm^2^ quantified at 10d and 28d post tamoxifen injection in WT^tdT^, HET^tdT^ and KO^tdT^ mice. **(G)** CSCs/mm^2^ quantified at 10d and 28d post tamoxifen injection in WT^tdT^, HET^tdT^ and KO^tdT^ mice. **(H)** CSCs/mm^2^ immunoreactive for pS6 quantified at 10d and 28d post tamoxifen injection for mice as in (**G**). **(I)** Representative images from contralateral TA muscle 10d post tamoxifen injection identifying Pax7+ QSCs in WT, HET and KO mice where arrowheads depict fluorescence as in (**D**). Scale bar: 20µm. **(J)** Representative images from contralateral TA muscle 28d post tamoxifen injection identifying tdT+ QSCs in WT^tdT^, HET^tdT^ and KO^tdT^ mice where arrowheads depict fluorescence as in (**D**). Scale bar: 20µm. For all statistical analysis, n = 3 mice per genotype, ordinary one-way ANOVA was used to compare groups. * P = < 0.05, ** P = < 0.003, *** P = 0.0003, **** P = < 0.0001.

Upon deleting TDP-43 pS6+ QSCs are preferentially lost compared to pS6-QSCs suggesting TDP-43 may be involved in transitioning SCs from quiescence to G_alert_To induce systemic G_Alert_ we injured one TA muscle and quantified SCs in the contralateral uninjured TA muscle (contralateral SCs or CSCs) at 10 and 28dpi (Fig. 6F-J). In WT mice, total SC abundance in was similar to that in uninjured muscle, but the number of pS6+ CSCs increased ∼2-fold at 10dpi, consistent with robust entry into G_Alert_ following a distant injury (Fig. 6G, H). In both HET and KO mice overall CSC numbers were reduced to ∼50% of WT but a striking deficit in G_Alert_ CSCs were observed. Only 20% of the total CSCs in contralateral muscle at 10dpi were pS6+ in HET and KO mice compared to ∼40% in WT mice (Fig. 6G-I). By 28dpi, WT^tdT^ mice pS6+ CSCs declined as the systemic response resolved (Fig. 6H,J). However, pS6+ CSCs were nearly undetectable in KO^tdT^ contralateral muscle at 28 dpi, with only ∼2 pS6+ CSCs per mm^2^ (Fig. 6H). In HET^tdT^ mice an intermediate phenotype of ∼6 pS6+ CSCs were observed, consistent with partial impairment of G_Alert_ entry when TDP-43 dosage is reduced (Fig. 6H).

G_alert_ pS6+ CSCs lacking TDP-43 are preferentially depleted in uninjured contralateral muscles following a distant injury and in injured TA muscles compared to overall SC loss. Therefore, TDP-43 appears required either for SC entry into G_Alert_ or for survival/maintenance within the G_alert_ state. Preferential loss of G_Alert_ SCs provides a mechanistic explanation for the progressive loss of SCs in uninjured muscle following TDP-43 ablation and supports the conclusion that TDP-43 sustains SC survival as the cells transit from quiescence into G_alert_.

## Discussion

Transitioning SCs from quiescence into the cell cycle during skeletal muscle maintenance and repair requires organelle biogenesis,^9,10^ cellular growth,^5^ metabolic remodeling,^5,32^ and increased cellular stress.^33,34^ Because post-transcriptional mechanisms regulate SC activation and cell cycle re-entry,^17–23^ and because TDP-43 loss prevents myogenesis,^25^ we asked whether TDP-43 regulates SC fate transitions. Ablating TDP-43 in SCs depleted SCs following a muscle injury eliminating muscle regenerative capacity. Moreover, TDP-43 loss in the absence of muscle injury progressively depleted the SC pool.

SCs exist in a niche between the basement membrane and the myofiber plasma membrane, responding to nearby and distant signals to maintain and regenerate muscle. Once SCs enter the cell cycle they either divide asymmetrically or symmetrically with daughter cells renewing the SC pool, proliferating, or differentiating to repair or rebuild skeletal muscle myofibers. By lineage labeling SCs with tdTomato we assessed whether SCs fuse into myofibers by quantifying tdTomato-labeled myofibers.^27^ In KO^tdT^ and HET^tdT^ mice, the number of tdTomato+ myofibers was significantly less than the numbers in WT^tdT^ controls, consistent with SC loss prior to fusion or impaired SC fusion.

TdTomato+ myofibers are unlikely to arise through vesicle-mediated transfer of tdTomato protein^35^ or tdTomato mRNA because WT^tdT^ and HET^tdT^ possessed indistinguishable SC numbers, yet tdTomato+ myofibers in HET^tdT^ were reduced greater than 2-fold compared to tdTomato+ myofibers in WT^tdT^ mice. While exosomal transfer of fluorescent proteins may occur, direct exosomal transfer is likely transient and rare since SCs comprise ∼3% or less of the cell number in skeletal muscle. Therefore, reducing or eliminating TDP-43 compromises the ability of SCs to repair or maintain myofibers in skeletal muscle.

Within the SC niche, quiescent SCs respond to nearby and distant signals for muscle maintenance and repair by entering a reversible G_Alert_ state.^5^ Following a distant muscle injury, 50-60% of SCs in contralateral uninjured muscle enter G_Alert_.^5^ G_Alert_ SCs are markedly reduced when TDP-43 is ablated, supporting the conclusion that TDP-43 is required for G_Alert_ entry or survival within G_Alert_ where TDP-43-null cells apoptose. Depleting G_Alert_ SCs explains the progressive loss of SCs in the absence of a muscle injury and the preferential loss of G_alert_ SCs from uninjured and injured muscle compared to overall SC loss. Rapidly depleting G_Alert_ SCs likely explains the progressive collapse of the SC pool and the failure of muscles to regenerate in TDP-43 conditional knockout mice.

To better understand mechanisms underlying TDP-43 function in SCs, we performed psuedotime trajectory analyses using single-cell RNA sequencing datasets identifying putative TDP-43-dependent stress response transcripts increasing as SCs transition from G_0_ to S-phase. Expression of these stress-associated transcripts is low in terminally differentiated myogenic cells and in mature myonuclei but highest as the myoblast progeny of SCs are expanding to generate new myonuclei. Cellular stress associated with quiescent SCs transitioning to rapidly proliferating progenitors likely induces cell stress responses for duplicating organelles, S-phase entry and mitosis to maintain proteostasis and DNA fidelity. TDP-43 may be involved in RNA processing of these cell stress transcripts where a decline or failure of these transcripts to generate proteins could compromise cell viability.

The requirement of TDP-43 for SC survival as SCs transit from quiescence to G_Alert_ has potentially broad implications. SC decline and regenerative failure in TDP-43 KO mice mirror aspects of muscle decline observed during aging and degenerative neuromuscular diseases associated with TDP-43 pathology.^36–40^ Given the prevalence of TDP-43 involvement in amyotrophic lateral sclerosis and inclusion body myositis, it is possible that impairing TDP-43 contributes to muscle degeneration in part through compromised SC maintenance.^36–40^ This idea is consistent with an increasing number of degenerative neuromuscular diseases arising from dynsfunctional SCs (satellite cell-opathies).^41,42^ Because loss of a single TDP-43 allele disrupts the transit of SCs into G_Alert_, or fails to maintain SCs in G_alert_ we believe partial reductions in SC TDP-43 protein impairs muscle maintenance. The precise TDP-43-dependent molecular mechanisms involved remain incompletely understood, and delineating these mechanisms may identify future strategies to preserve SC function in degenerative neuromuscular diseases with an SC involvement.

## Resource Availability

### Lead Contact

For reagents, resource requests, and further information, please contact Bradley Olwin.

### Data and materials availability

All data are available in the main text or the supplementary materials. Enhanced CLIP data is available on GEO (https://www.ncbi.nlm.nih.gov/geo/query/acc.cgi?acc=GSE104796) under accession number GEO: GSE104796 and sequencing data have been deposited on GEO under accession number GEO: GSE180225. All mice and cell lines generated from mice in this study are available upon request from the lead contact with a completed Materials Transfer Agreement.

## Materials and Methods

### Experimental design

Our goal in this study is to investigate the functional role of TDP-43 in satellite cells using inducible KO mice and cell lines as described below.

### Mice

Mice were bred and housed according to National Institute of Health (NIH) guidelines for the ethical treatment of animals in a pathogen-free facility at the University of Colorado at Boulder. The University of Colorado Institutional Animal Care and Use Committee (IACUC) approved all animal protocols and procedures to ensure all studies complied with all ethical regulations. Mice used were either C57BL6 (Jackson Labs Stock No. 000664), *Pax7^CreERT2^* (017763), *Tardbp^loxp/loxp^*^25,43^ (017589), or *Rosa26^tdTomato^*(007914) mice. Crossing *Tardbp^loxp/loxp^* or *Rosa26^tdTomato^*mice into *Pax7^CreERT2^* generated conditional *Tardbp^loxp/+^* or *Tardbp^loxp/loxp^* mice. Crossing conditional *Tardbp^loxp/+^* or *Tardbp^loxp/loxp^* mice with *Rosa26^tdTomato^* mice generated *Tardbp^loxp/+^*; *Rosa26^tdTomato^*or *Tardbp^loxp/loxp^*;*Rosa26^tdTomato^* mice, respectively. TA muscles were isolated from mice between 6 and 8 months old and were a mix of male and female. Control mice were randomly assigned and were sex and age matched to the mice and crosses outlined above. Sample sizes were set at *n* = 3. Each mouse used in this study was genotyped by Transnetyx (Cordova, TN).

### Mouse injuries and Tamoxifen

Mice were anesthetized with isoflurane, TA muscles injected with 50µL of 1.2% BaCl_2_ in sterile saline injuring the extensor digitorious longus and the TA muscle.^44^ Tamoxifen (Sigma-Aldrich) was resuspended in corn oil (Sigma-Aldrich) and administered by injecting 0.075 mg of tamoxifen per gram of mouse weight intraperitoneally. All muscle injuries were made blinded to genotype.

### Immunodetection of antigens in tissue sections

TA muscles were dissected, fixed on ice for 2 h with 4% paraformaldehyde, and transferred to phosphate-buffered saline (PBS) with 30% sucrose at 4°C overnight. Muscles were mounted in OCT (Tissue-Tek), cryo-sectioning was performed on a Leica cryostat to generate 10-μm thick sections, and sections stored at −80°C until staining. Tissue sections were post-fixed in 4% paraformaldehyde for 10 min at room temperature and washed three times for 5 min in PBS. Immunodetection of PAX7, TDP-43, cytochrome c, cleaved PARP1, and pS6 required heat-induced epitope antigen retrieval, for which post-fixed slides were placed in citrate buffer, pH 6.0, and subjected to 6 min of high pressure-cooking in a Cuisinart model CPC-600 pressure cooker. For immunodetection, tissue sections were permeabilized with 0.25% Triton X-100 (Sigma-Aldrich) in PBS containing 2% bovine serum albumin (BSA) (Sigma-Aldrich) for 60 min at room temperature following by incubation with a primary antibody at 4 °C overnight and by incubation with a secondary antibody at room temperature for 1 h. Primary antibodies included mouse anti-PAX7 (Developmental Studies Hybridoma Bank) at 1:750, rabbit anti-TDP-43 (ProteinTech) at 1:200, mouse anti-TDP-43 (Abcam) at 1:200, mouse anti-cytochrome C at 1:200, mouse anti-cleaved PARP1 at 1:200, and rabbit anti-pS6 (Cell Signaling Technologies) at 1:200.Alexa secondary antibodies included donkey anti-rabbit 488 and 647, goat anti-mouse IgG2b 488, goat anti-mouse IgG2a 647, and goat anti-mouse IgG1 488, 555, and 647 (Thermo) were used at a dilution of 1:1,000. Wheat germ agglutinin Oregon Green 488 (Thermo) was used at 1:200 for visualization of myofiber membranes. Sections were incubated with 1 μg ml−1 DAPI for 10 min at room temperature then mounted in Mowiol supplemented with DABCO (Sigma-Aldrich) or ProLong Gold (Thermo) as an anti-fade agent.

### Picrosirius Red Staining

TA sections were prepared as described and stored at −80°C until staining was performed. Tissue was post-fixed in Bouin’s solution (Sigma-Aldrich) for 1 h at 56°C. Slides were then rinsed under tap water until fixative was removed and Picrosirius Red (American Master Tech) was added for 1 h at room temperature. Slides were then rinsed twice in 0.5% acetic acid (J.T. Baker) for one minute each, and tissue sections dehydrated step-wise in 50%, 75%, and 100% ethanol (Sigma-Aldrich) for 30 seconds each. Slides were then rinsed twice in xylene (VWR) for 1 minute and dried at room temperature before mounting with permount (ThermoFisher Scientific).

### Immunodetection of antigens in cultured cells

Cultured cells were washed with PBS in a laminar flow hood and fixed with 4% paraformaldehyde for 10 min at room temperature in a chemical hood. Cells were permeabilized with 0.25% Triton X-100 in PBS containing 2% BSA (Sigma-Aldrich) for 1 h at room temperature, incubated with primary antibody at 4 °C overnight followed by incubating with secondary antibodies at room temperature for 1 h. Primary antibodies included mouse anti-PAX7 (Developmental Studies Hybridoma Bank) at 1:750, rabbit anti-TDP-43 (ProteinTech) at 1:200, and mouse anti-TDP-43 (Abcam) at 1:200. Alexa secondary antibodies (ThermoFisher Scientific) were used at a dilution of 1:1,000. All antibodies were diluted in 0.125% Triton X-100 in PBS containing 2% BSA. Cells were incubated with 1 μg ml−1 DAPI for 10 min at room temperature then mounted in Mowiol supplemented with DABCO (Sigma-Aldrich) as an anti-fade agent prior to imaging.

### Microscopy and image analysis

Processed tissue and cells were imaged on a Nikon inverted spinning disk confocal microscope where objectives used were 10x/0.45NA Plan Apo, 20x/0.75NA Plan Apo, and 60x/1.4NA Plan Apo. For imaging of of Picrosirus red stains, a Nikon TiU widefield with RGB color brightfield was used with a 4x 0.13NA Plan Fluor WD objective. All images were processed using Fiji ImageJ. Confocal stacks were projected as maximum intensity projections for each channel and merged to form a single image. Brightness and contrast were adjusted for the entire image when necessary.

### Isolation of primary muscle stem cells

Gastrocnemius, extensor digitorum longus, tibialis anterior and all other lower hindlimb muscles were dissected from mice and placed into 400U/mL collagenase (Worthington) at 37°C for 1 h with intermittent vortexing for 1 min at medium speed every 10 min. Collagenase was quenched by adding Ham’s F-12C supplemented with 15% horse serum (Gibco). Cells were filtered through three strainers of 100 µm, 70 µm, and 40μm (BD Falcon) each; the final flow through was centrifuged at 1500×g for 5 min and the cell pellets re-suspended in Ham’s F-12C supplemented with 15% horse serum and FGF-2 (50 nM). Isolated cells were then either harvested immediately for RNA isolation (Qiagen) or plated on gelatin-coated 15-cm tissue-culture dishes (Corning) for culture.

### TDP-43cKO cell line immortalization and culture

Primary SCs from *Pax7^CreERT2^*;*Tardbp^loxp/loxp^*;*Rosa26^tdTomato^*mice were spontaneously immortalized as described elsewhere.^45–47^ Primary SCs were isolated as and plated at clonal density on gelatin coated dishes in Ham’s F-12C supplemented with penicillin and streptomycin, 15% horse serum (Gibco), 20% fetal bovine serum (Sigma-Aldrich), and FGF-2 (50 nM). Cells were allowed to grow and establish individual colonies from a single cell. Cultured clones were rinsed with sterile PBS and isolated using glass cloning rings with 0.25% trypsin-EDTA (Sigma-Aldrich). Clones were serial passaged until a uniformly proliferating cell line was achieved and their ability to differentiate to form myotubes. For conditional deletion of TDP-43, cells 4-hydroxytamoxifen (400ng/mL, Sigma-Aldrich) resuspended in 100% molecular biology grade ethanol (Sigma-Aldrich) was added and control cells were treated with EtOH vehicle.

### Flow Cytometry

TDP-43cKO cells were grown in the presence or absence of EtOH (vehicle control), 400ng/mL 4-hydroxytamoxifen, or 1uM staurosporin (Sigma-Aldrich) for 72 h. Prior to analysis, cells were gently washed with PBS and removed from the dish with trypsin, pelleted, and resuspended in buffer with Annexin V to identify apoptotic cells and DAPI to identify necrotic cells or cells with compromised membrane integrity (Elabscience). Cells were analyzed on a Beckman Coulter CyAn ADP Analyzer.

### Live Cell Microscopy

Immortalized TDP-43cKO cells were grown in the presence or absence of 4-hydroxytamoxifen as described above on gelatin coated glass-bottom imaging dishes (MatTek) and cell death was detected with ReadyProbes Cell Viability Imaging Kit (Life Technologies). Live cell microscopy was performed by timelapse imaging, where cells were imaged every 10 minutes on an Olympus CellVoyager CV1000 spinning disk using a 4x 0.13NA UPlanFl objective. Timelapse videos were processed and analyzed using Fiji ImageJ.

### TDP-43 eClip and RNA sequencing

Genes with −log10pvalues greater than 5 were categorized with Panther^48–50^ for Mus musculus and the Panther-GO biological process complete annotation set from a prior published TDP43-eCLIP dataset.^25^ The top 10 GO terms based on fold-enrichment were identified and GO biological process complete terms visually organized by similarity with Revigo into multidimensional maps and modified with R.^51^ Circle size indicates uniqueness, which identifies indispensable terms compared to the whole list, and the color range reflects the Panther-GO adjusted p values. Points were labeled if the Panther-GO p value was less than 10^−7^ for the myoblast dataset (low of −10) and less than 10^−11^ for the myotube dataset (low of - 20).

### Q-PCR

#### Immortalized TDP-43cKO cell line qPCR analysis

TDP-43 KO cells (passage 13) were plated 8.5k cells per 6-well and treated with 400ng/mL 4-Hydroxytamoxifen or EtOH vehicle control for 72hrs to induce TDP-43 knock-out in cells expressing Pax7. RNA was isolated using Qiagen RNeasy minikit (Cat. 74104) and cDNA synthesis and subsequent qPCR performed using the Superscript IV First-strand synthesis system (Cat. 18091050). QPCR values were assessed by ΔΔCT analysis and normalized to Pax7 expression. Statistical analysis was performed using 2-way Anova in GaphPad Prism (v. 10).

### Single-nuclear RNAseq

A subset of the prior published data for single-nuclear RNAseq,^31^ the myogenic cell populations from adult mice,were analyzed with Seurat v5.1.0.^52^ The resulting Seurat object was passed through FindVariableFeatures(), ScaleData(), RunPCA(), FindNeighbors(), FindClusters(), and RunUMAP() with dims of 1:3 and n.neighbors of 25L. For the pseudotime analysis, we used Monocle3 v1.3.5^53–55^ on the Seurat object to generate trajectories with the uninjured MuSC population manually set as the root module.

### Statistical Analysis

Statistical analyses were conducted using unpaired *t* tests or analysis of variance (ANOVA). A confidence interval of 95% was used. Statistical analyses were performed with GraphPad Prism (v. 10).

## Acknowledgments

We thank James Orth and acknowledge the Light Microscopy Core Facility, Porter Biosciences B047, B049, B051, and B059 at the University of Colorado Boulder (RRID:SCR_018993) for help and advice with microscopy; Nicole Dalla Betta for help with immunofluorescence and tissue sectioning; and Alicia Cutler for intellectual input.

## Author contributions

Conceptualization: TEE, BBO

Methodology: TEE, BBO

Investigation: TEE, SD, TKV, HC, TE

Visualization: TEE, SD, TKV, HC

Supervision: BBO

Writing—original draft: TEE, BBO

Writing—review & editing: TEE, BBO

## References

1. Murphy, M.M., Lawson, J.A., Mathew, S.J., Hutcheson, D.A., and Kardon, G. (2011). Satellite cells, connective tissue fibroblasts and their interactions are crucial for muscle regeneration. Development 138, 3625–3637. 10.1242/dev.064162.

2. Lepper, C., Partridge, T.A., and Fan, C.-M. (2011). An absolute requirement for Pax7-positive satellite cells in acute injury-induced skeletal muscle regeneration. Development 138, 3639–3646. 10.1242/dev.067595.

3. Sambasivan, R., Yao, R., Kissenpfennig, A., Van Wittenberghe, L., Paldi, A., Gayraud-Morel, B., Guenou, H., Malissen, B., Tajbakhsh, S., and Galy, A. (2011). Pax7-expressing satellite cells are indispensable for adult skeletal muscle regeneration. Development 138, 3647–3656. 10.1242/dev.067587.

4. Dumont, N.A., Bentzinger, C.F., Sincennes, M.-C., and Rudnicki, M.A. (2015). Satellite Cells and Skeletal Muscle Regeneration. In Comprehensive Physiology (John Wiley & Sons, Ltd), pp. 1027–1059. 10.1002/cphy.c140068.

5. Rodgers, J.T., King, K.Y., Brett, J.O., Cromie, M.J., Charville, G.W., Maguire, K.K., Brunson, C., Mastey, N., Liu, L., Tsai, C.-R., et al. (2014). mTORC1 controls the adaptive transition of quiescent stem cells from G0 to GAlert. Nature 510, 393–396. 10.1038/nature13255.

6. Zhang, P., Liang, X., Shan, T., Jiang, Q., Deng, C., Zheng, R., and Kuang, S. (2015). mTOR is necessary for proper satellite cell activity and skeletal muscle regeneration. Biochem. Biophys. Res. Commun. 463, 102–108. 10.1016/j.bbrc.2015.05.032.

7. Yue, F., Bi, P., Wang, C., Li, J., Liu, X., and Kuang, S. (2016). Conditional Loss of *Pten* in Myogenic Progenitors Leads to Postnatal Skeletal Muscle Hypertrophy but Age-Dependent Exhaustion of Satellite Cells. Cell Rep. 17, 2340–2353. 10.1016/j.celrep.2016.11.002.

8. Brun, C.E., Sincennes, M.-C., Lin, A.Y.T., Hall, D., Jarassier, W., Feige, P., Le Grand, F., and Rudnicki, M.A. (2022). GLI3 regulates muscle stem cell entry into GAlert and self-renewal. Nat. Commun. 13, 3961. 10.1038/s41467-022-31695-5.

9. Wozniak, A.C., Kong, J., Bock, E., Pilipowicz, O., and Anderson, J.E. (2005). Signaling satellite-cell activation in skeletal muscle: Markers, models, stretch, and potential alternate pathways. Muscle Nerve 31, 283–300. 10.1002/mus.20263.

10. Anderson, J.E. (2000). A Role for Nitric Oxide in Muscle Repair: Nitric Oxide–mediated Activation of Muscle Satellite Cells. Mol. Biol. Cell 11, 1859–1874. 10.1091/mbc.11.5.1859.

11. Troy, A., Cadwallader, A.B., Fedorov, Y., Tyner, K., Tanaka, K.K., and Olwin, B.B. (2012). Coordination of satellite cell activation and self-renewal by Par-complex-dependent asymmetric activation of p38alpha/beta MAPK. Cell Stem Cell 11, 541–553. 10.1016/j.stem.2012.05.025.

12. Jones, N.C., Tyner, K.J., Nibarger, L., Stanley, H.M., Cornelison, D.D.W., Fedorov, Y.V., and Olwin, B.B. (2005). The p38α/β MAPK functions as a molecular switch to activate the quiescent satellite cell. J. Cell Biol. 169, 105–116. 10.1083/jcb.200408066.

13. Zammit, P.S. (2017). Function of the myogenic regulatory factors Myf5, MyoD, Myogenin and MRF4 in skeletal muscle, satellite cells and regenerative myogenesis. Semin. Cell Dev. Biol. 72, 19–32. 10.1016/j.semcdb.2017.11.011.

14. Sincennes, M.-C., Brun, C.E., and Rudnicki, M.A. (2016). Concise Review: Epigenetic Regulation of Myogenesis in Health and Disease. Stem Cells Transl. Med. 5, 282–290. 10.5966/sctm.2015-0266.

15. Farina, N.H., Hausburg, M., Betta, N.D., Pulliam, C., Srivastava, D., Cornelison, D., and Olwin, B.B. (2012). A role for RNA post-transcriptional regulation in satellite cell activation. Skelet Muscle 2, 21. 10.1186/2044-5040-2-21.

16. Cho, D.S., and Doles, J.D. (2017). Single cell transcriptome analysis of muscle satellite cells reveals widespread transcriptional heterogeneity. Gene 636, 54–63. 10.1016/j.gene.2017.09.014.

17. Hausburg, M.A., Doles, J.D., Clement, S.L., Cadwallader, A.B., Hall, M.N., Blackshear, P.J., Lykke-Andersen, J., and Olwin, B.B. (2015). Post-transcriptional regulation of satellite cell quiescence by TTP-mediated mRNA decay. Elife 4, e03390. 10.7554/eLife.03390.

18. Morrée, A. de, Velthoven, C.T.J. van, Gan, Q., Salvi, J.S., Klein, J.D.D., Akimenko, I., Quarta, M., Biressi, S., and Rando, T.A. (2017). Staufen1 inhibits MyoD translation to actively maintain muscle stem cell quiescence. Proc. Natl. Acad. Sci. 114, E8996–E9005. 10.1073/pnas.1708725114.

19. Yue, L., Wan, R., Luan, S., Zeng, W., and Cheung, T.H. (2020). Dek Modulates Global Intron Retention during Muscle Stem Cells Quiescence Exit. Dev. Cell 53, 661–676.e6. 10.1016/j.devcel.2020.05.006.

20. Cheung, T.H., Quach, N.L., Charville, G.W., Liu, L., Park, L., Edalati, A., Yoo, B., Hoang, P., and Rando, T.A. (2012). Maintenance of muscle stem-cell quiescence by microRNA-489. Nature 482, 524–528. 10.1038/nature10834.

21. Zismanov, V., Chichkov, V., Colangelo, V., Jamet, S., Wang, S., Syme, A., Koromilas, A.E., and Crist, C. (2015). Phosphorylation of eIF2alpha Is a Translational Control Mechanism Regulating Muscle Stem Cell Quiescence and Self-Renewal. Cell Stem Cell. 10.1016/j.stem.2015.09.020.

22. Fujita, R., Lean, G., Jamet, S., Hébert, S., Kleinman, C.L., and Crist, C. (2020). Satellite cell expansion is mediated by P-eIF2α dependent *Tacc3* translation (Cell Biology) 10.1101/2020.05.13.093302.

23. Chenette, D.M., Cadwallader, A.B., Antwine, T.L., Larkin, L.C., Wang, J., Olwin, B.B., and Schneider, R.J. (2016). Targeted mRNA Decay by RNA Binding Protein AUF1 Regulates Adult Muscle Stem Cell Fate, Promoting Skeletal Muscle Integrity. Cell Rep. 16, 1379–1390. 10.1016/j.celrep.2016.06.095.

24. Wheeler, J.R., Whitney, O.N., Vogler, T.O., Nguyen, E.D., Pawlikowski, B., Lester, E., Cutler, A., Elston, T., Dalla Betta, N., Parker, K.R., et al. (2022). RNA-binding proteins direct myogenic cell fate decisions. eLife 11, e75844. 10.7554/eLife.75844.

25. Vogler, T.O., Wheeler, J.R., Nguyen, E.D., Hughes, M.P., Britson, K.A., Lester, E., Rao, B., Betta, N.D., Whitney, O.N., Ewachiw, T.E., et al. (2018). TDP-43 and RNA form amyloid-like myo-granules in regenerating muscle. Nature 563, 508–513. 10.1038/s41586-018-0665-2.

26. Murach, K.A., Vechetti, I.J., Van Pelt, D.W., Crow, S.E., Dungan, C.M., Figueiredo, V.C., Kosmac, K., Fu, X., Richards, C.I., Fry, C.S., et al. (2020). Fusion-Independent Satellite Cell Communication to Muscle Fibers During Load-Induced Hypertrophy. Function 1, zqaa009. 10.1093/function/zqaa009.

27. Pawlikowski, B., Pulliam, C., Betta, N.D., Kardon, G., and Olwin, B.B. (2015). Pervasive satellite cell contribution to uninjured adult muscle fibers. Skelet. Muscle 5, 42. 10.1186/s13395-015-0067-1.

28. Ji, S., Ma, P., Cao, X., Wang, J., Yu, X., Luo, X., Lu, J., Hou, W., Zhang, Z., Yan, Y., et al. (2022). Myoblast-derived exosomes promote the repair and regeneration of injured skeletal muscle in mice. FEBS Open Bio 12, 2213–2226. 10.1002/2211-5463.13504.

29. Fry, C.S., Kirby, T.J., Kosmac, K., McCarthy, J.J., and Peterson, C.A. (2017). Myogenic Progenitor Cells Control Extracellular Matrix Production by Fibroblasts during Skeletal Muscle Hypertrophy. Cell Stem Cell 20, 56–69. 10.1016/j.stem.2016.09.010.

30. Bernet, J.D., Doles, J.D., Hall, J.K., Kelly Tanaka, K., Carter, T.A., and Olwin, B.B. (2014). p38 MAPK signaling underlies a cell-autonomous loss of stem cell self-renewal in skeletal muscle of aged mice. Nat Med 20, 265–271. 10.1038/nm.3465.

31. Kurland, J.V., Cutler, A.A., Stanley, J.T., Betta, N.D., Deusen, A.V., Pawlikowski, B., Hall, M., Antwine, T., Russell, A., Allen, M.A., et al. (2023). Aging disrupts gene expression timing during muscle regeneration. Stem Cell Rep. 18, 1325–1339. 10.1016/j.stemcr.2023.05.005.

32. Sousa-Victor, P., García-Prat, L., and Muñoz-Cánoves, P. (2022). Control of satellite cell function in muscle regeneration and its disruption in ageing. Nat. Rev. Mol. Cell Biol. 23, 204–226. 10.1038/s41580-021-00421-2.

33. Harding, H.P., Zhang, Y., Zeng, H., Novoa, I., Lu, P.D., Calfon, M., Sadri, N., Yun, C., Popko, B., Paules, R., et al. (2003). An Integrated Stress Response Regulates Amino Acid Metabolism and Resistance to Oxidative Stress. Mol. Cell 11, 619–633. 10.1016/S1097-2765(03)00105-9.

34. Machado, L., Geara, P., Camps, J., Dos Santos, M., Teixeira-Clerc, F., Van Herck, J., Varet, H., Legendre, R., Pawlotsky, J.-M., Sampaolesi, M., et al. (2021). Tissue damage induces a conserved stress response that initiates quiescent muscle stem cell activation. Cell Stem Cell 28, 1125–1135.e7. 10.1016/j.stem.2021.01.017.

35. Vechetti Jr, I.J., Valentino, T., Mobley, C.B., and McCarthy, J.J. (2021). The role of extracellular vesicles in skeletal muscle and systematic adaptation to exercise. J. Physiol. 599, 845–861. 10.1113/JP278929.

36. Mori, F., Tada, M., Kon, T., Miki, Y., Tanji, K., Kurotaki, H., Tomiyama, M., Ishihara, T., Onodera, O., Kakita, A., et al. (2019). Phosphorylated TDP-43 aggregates in skeletal and cardiac muscle are a marker of myogenic degeneration in amyotrophic lateral sclerosis and various conditions. Acta Neuropathol. Commun. 7, 165. 10.1186/s40478-019-0824-1.

37. D’Agostino, C., Nogalska, A., Engel, W.K., and Askanas, V. (2011). In sporadic inclusion body myositis muscle fibres TDP-43-positive inclusions are less frequent and robust than p62 inclusions, and are not associated with paired helical filaments. Neuropathol. Appl. Neurobiol. 37, 315–320. 10.1111/j.1365-2990.2010.01108.x.

38. Arai, T., Hasegawa, M., Akiyama, H., Ikeda, K., Nonaka, T., Mori, H., Mann, D., Tsuchiya, K., Yoshida, M., Hashizume, Y., et al. (2006). TDP-43 is a component of ubiquitin-positive tau-negative inclusions in frontotemporal lobar degeneration and amyotrophic lateral sclerosis. Biochem. Biophys. Res. Commun. 351, 602–611. 10.1016/j.bbrc.2006.10.093.

39. Wilson, R.S., Yu, L., Trojanowski, J.Q., Chen, E.-Y., Boyle, P.A., Bennett, D.A., and Schneider, J.A. (2013). TDP-43 Pathology, Cognitive Decline, and Dementia in Old Age. JAMA Neurol. 70, 1418–1424. 10.1001/jamaneurol.2013.3961.

40. Sorarú, G., Orsetti, V., Buratti, E., Baralle, F., Cima, V., Volpe, M., D’ascenzo, C., Palmieri, A., Koutsikos, K., Pegoraro, E., et al. (2010). TDP-43 in skeletal muscle of patients affected with amyotrophic lateral sclerosis. Amyotroph. Lateral Scler. 11, 240–243. 10.3109/17482960902810890.

41. Ganassi, M., Muntoni, F., and Zammit, P.S. (2022). Defining and identifying satellite cell-opathies within muscular dystrophies and myopathies. Exp. Cell Res. 411, 112906. 10.1016/j.yexcr.2021.112906.

42. Ganassi, M., and Zammit, P.S. (2022). Involvement of muscle satellite cell dysfunction in neuromuscular disorders: Expanding the portfolio of satellite cell-opathies. Eur. J. Transl. Myol. 32. 10.4081/ejtm.2022.10064.

43. Chiang, P.-M., Ling, J., Jeong, Y.H., Price, D.L., Aja, S.M., and Wong, P.C. (2010). Deletion of TDP-43 down-regulates Tbc1d1, a gene linked to obesity, and alters body fat metabolism. Proc. Natl. Acad. Sci. 107, 16320–16324. 10.1073/pnas.1002176107.

44. Morton, A.B., Norton, C.E., Jacobsen, N.L., Fernando, C.A., Cornelison, D.D.W., and Segal, S.S. (2019). Barium chloride injures myofibers through calcium-induced proteolysis with fragmentation of motor nerves and microvessels. Skelet. Muscle 9, 27. 10.1186/s13395-019-0213-2.

45. Hauschka, S.D., Linkhart, T.A., Clegg, C., and Merrill, G. (1979). Clonal studies of human and mouse muscle. Muscle Regen., 311–322.

46. David Yaffe and Ora Saxel (1977). Serial passaging and differentiation of myogenic cells isolated from dystrophic mouse muscle. Nature 270, 725–727.

47. Nowak, J.A., Malowitz, J., Girgenrath, M., Kostek, C.A., Kravetz, A.J., Dominov, J.A., and Miller, J.B. (2004). Immortalization of mouse myogenic cells can occur without loss of p16INK4a, p19ARF, or p53 and is accelerated by inactivation of Bax. BMC Cell Biol. 5, 1. 10.1186/1471-2121-5-1.

48. Thomas, P.D., Ebert, D., Muruganujan, A., Mushayahama, T., Albou, L.-P., and Mi, H. (2022). PANTHER: Making genome-scale phylogenetics accessible to all. Protein Sci. 31, 8–22. 10.1002/pro.4218.

49. Mi, H., Muruganujan, A., Huang, X., Ebert, D., Mills, C., Guo, X., and Thomas, P.D. (2019). Protocol Update for large-scale genome and gene function analysis with the PANTHER classification system (v.14.0). Nat. Protoc. 14, 703–721. 10.1038/s41596-019-0128-8.

50. Mi, H., and Thomas, P. (2009). PANTHER Pathway: an ontology-based pathway database coupled with data analysis tools. Methods Mol. Biol. Clifton NJ 563, 123–140. 10.1007/978-1-60761-175-2_7.

51. Supek, F., Bošnjak, M., Škunca, N., and Šmuc, T. (2011). REVIGO Summarizes and Visualizes Long Lists of Gene Ontology Terms. PLOS ONE 6, e21800. 10.1371/journal.pone.0021800.

52. Hao, Y., Stuart, T., Kowalski, M.H., Choudhary, S., Hoffman, P., Hartman, A., Srivastava, A., Molla, G., Madad, S., Fernandez-Granda, C., et al. (2024). Dictionary learning for integrative, multimodal and scalable single-cell analysis. Nat. Biotechnol. 42, 293–304. 10.1038/s41587-023-01767-y.

53. Trapnell, C., Cacchiarelli, D., Grimsby, J., Pokharel, P., Li, S., Morse, M., Lennon, N.J., Livak, K.J., Mikkelsen, T.S., and Rinn, J.L. (2014). The dynamics and regulators of cell fate decisions are revealed by pseudotemporal ordering of single cells. Nat. Biotechnol. 32, 381–386. 10.1038/nbt.2859.

54. Qiu, X., Mao, Q., Tang, Y., Wang, L., Chawla, R., Pliner, H., and Trapnell, C. (2017). Reversed graph embedding resolves complex single-cell developmental trajectories. Genomics.

55. Cao, J., Spielmann, M., Qiu, X., Huang, X., Ibrahim, D.M., Hill, A.J., Zhang, F., Mundlos, S., Christiansen, L., Steemers, F.J., et al. (2019). The single-cell transcriptional landscape of mammalian organogenesis. Nature 566, 496–502. 10.1038/s41586-019-0969-x.

