## Supplemental for "TDP-43 Sustains Satellite Cells to Maintain and Regenerate Skeletal Muscle"

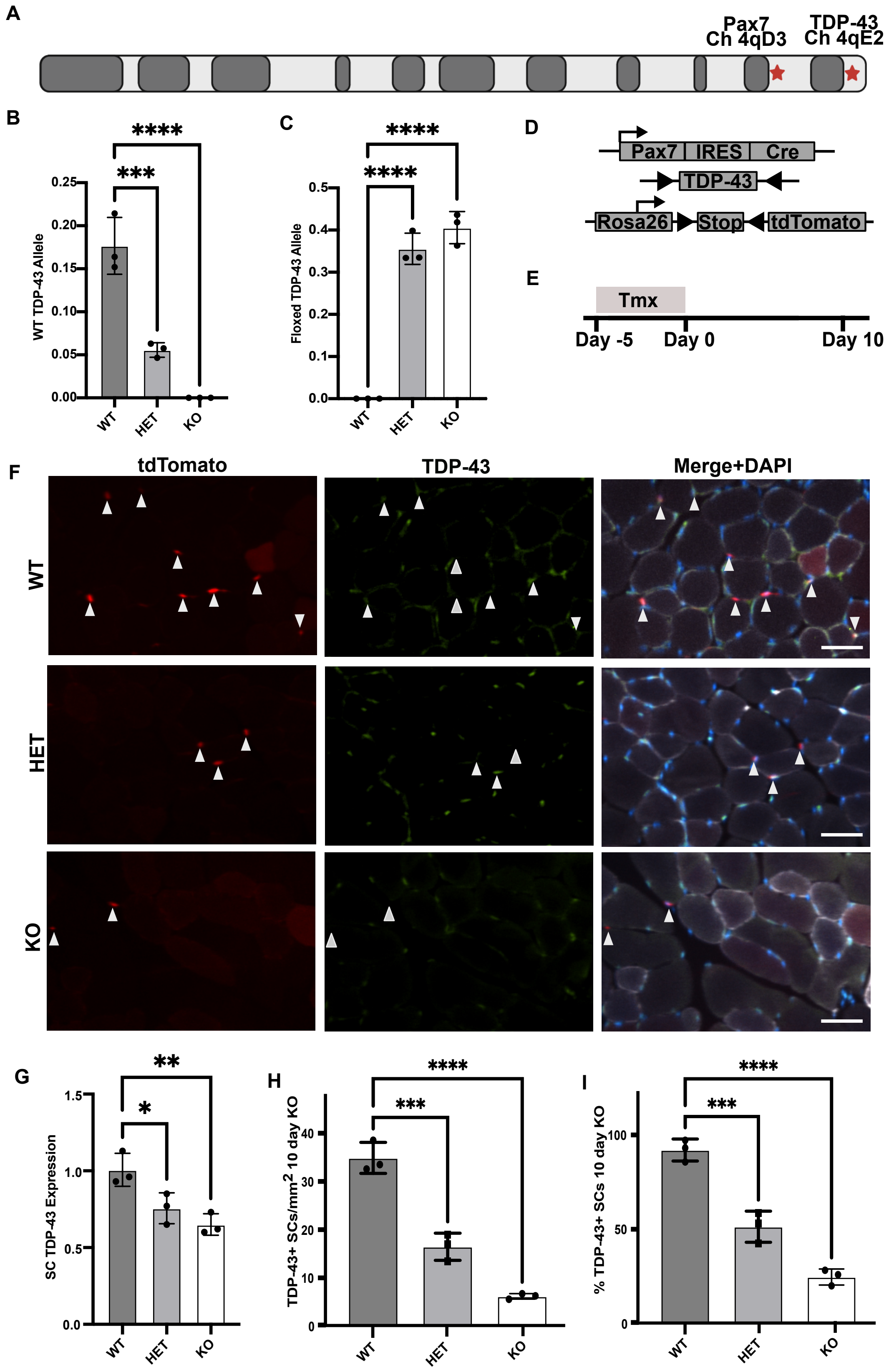


**Fig. S1: SCs require TDP-43 to Regenerate Muscle. (A)** Conditional TDP-43 KO mouse ideogram depicting proximity of TDP-43 and Pax7 loci on chromosome 4. **(B)** Genotyping from mouse tail biopsies of TDP-43cKO mice for WT TDP-43

gene using real-time PCR. **(C)** Genotyping of TDP-43cKO mice for floxed TDP-43 allele. **(D)** Schematic depicting conditional TDP-43 KO mice with addition of tdTomato, located at the ROSA 26 locus, allowing for detection of SCs and any myofibers that have undergone SC fusion. **(E)** Timeline for tamoxifen injection and time of injury and collection for TA muscles. **(F)** Representative images of TA muscle sections from $Pax7^{CreERT2}$;$TDP\text{-}43^{flox/flox}$;LSL:tdTomato (WT), $Pax7^{CreERT2}$;$TDP\text{-}43^{flox/wt}$;LSL:tdTomato (HET), and $Pax7^{CreERT2}$;$TDP\text{-}43^{flox/flox}$;LSL:tdTomato (WT) mice 10d following tamoxifen injection. Recombined SCs are tdTomato+ and TDP-43 visualized by fluorescent immunoreactivity. Solid arrowheads identify tdTomato +/TDP-43+ SCs. Scale bar = 50$\mu$m. **(G)** qRT-PCR for TDP-43 in SCs purified from 10d WT, HET and KO TA muscles. **(H)** TDP-43-expressing SCs/$mm^2$ 10d post tamoxifen injection for TA muscles from WT, HET and KO mice. **(I)** Percent TDP-43 positive SCs at 10d post tamoxifen injection for TA muscles from WT, HET and KO mice. For B, C, G-I n = 3 mice per genotype. For all statistical analysis, Ordinary one-way ANOVA was used to compare groups. * $P = < 0.05$, ** $P = < 0.007$, *** $P = < 0.0005$, **** $P = < 0.0001$.

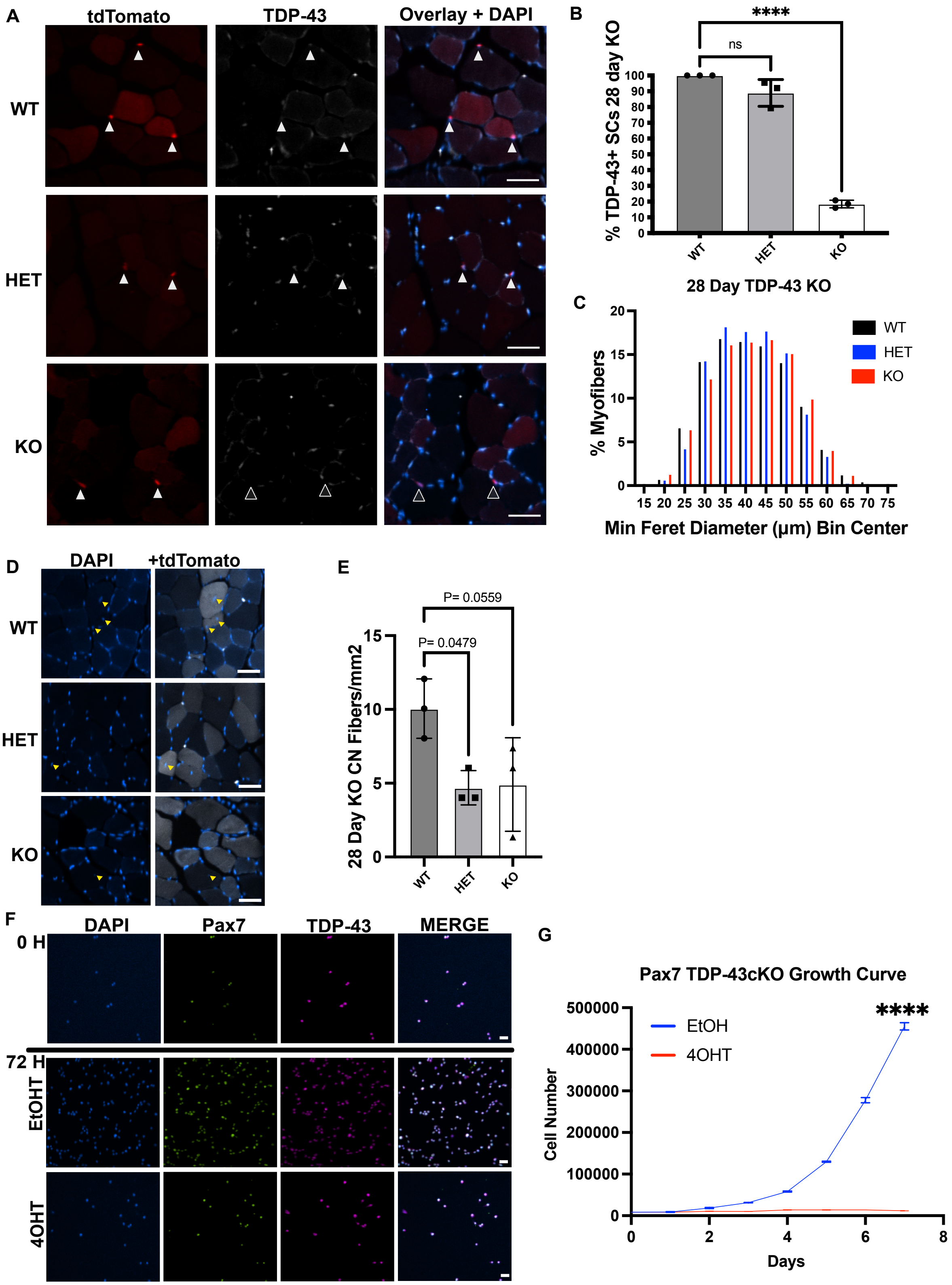


**Fig. S2. TDP-43 null SCs are unable to maintain muscle**. **(A)** Representative TA muscle cross-section images from $WT^{tdT}$, $HET^{tdT}$ and KOtdT mice 28-day following tamoxifen injections. TdTomato+ SCs and TDP-43+ SCs are indicated by solid arrowheads with outlined arrowheads identifying TDP-43- SCs. Scale bar:

50$\mu$m **(B)** Percent SCs expressing TDP-43 in mice as in (**A**) 28d post tamoxifen injections. n = 3 mice. One-way ANOVA **** P = < 0.0001 **(C)** Frequency distribution of myofiber diameter in mice as in (**A**) 28d post tamoxifen injections. One-way ANOVA. Not significant. **(D)** Representative TA muscle cross-section images from mice in (A) 28 d post tamoxifen injections identifying centrally located nuclei. Arrowheads indicate centrally located nuclei. **(E)** Cross sections of TA in (D) were quantified for centrally located nuclei. One-way ANOVA P = < 0.05. **(F)** Representative images of recombined and unrecombined KO cells following EtOH or 4OHT treatment at time of cell plating (t = 0h) and at 72 h post-treatment. Scale bar = 25$\mu$m. **(G)** Growth curve plotted for cells treated as in (**H**). n = 3 independent biological experiments. Two-way ANOVA. **** P = < 0.0001.

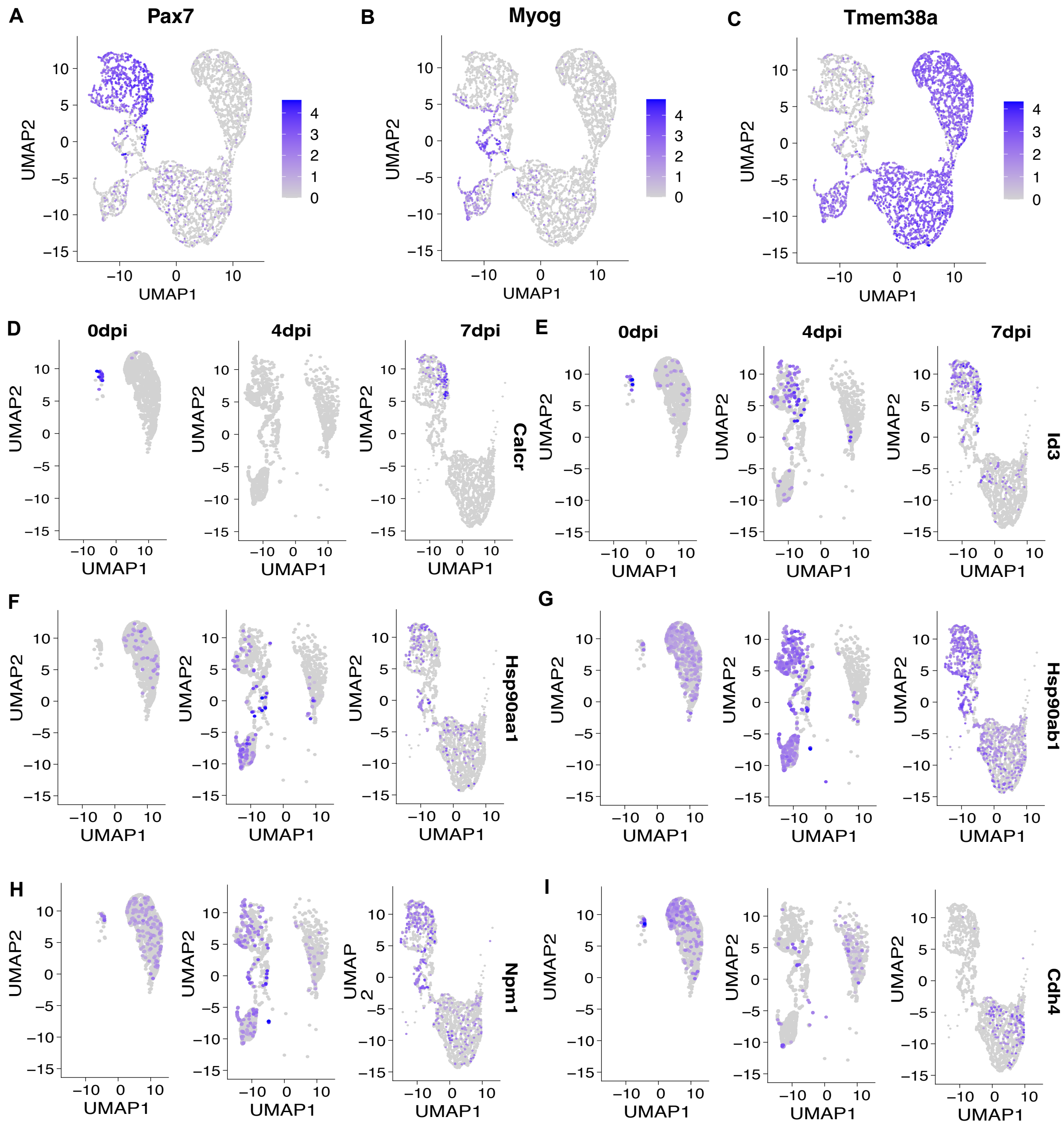


**Fig. S5. TDP-43-dependent stress response transcripts are regulated during cell fate changes. (A-C)** UMAP of SC clusters identified by Pax7 expression (A), progenitors by Myog expression (B) and myonuclei by Tmem38a expression (C). **(D-I)** UMAP clustering of nuclei from uninjured muscle and regenerating muscle depicting a time course for expression of (**D, E**) myogenic cell identifiers (Caclr and Id3) and (**F-I**) the TDP-43 cell-stress target genes (Hsp90aa1, HSP90ab1, Npm1, Cdh4) in uninjured muscle at 4 dpi and 7dpi.
